# Engineered and decellularized human cartilage graft exhibits intrinsic immuno-evasive properties and full skeletal repair capacity

**DOI:** 10.1101/2025.01.10.632347

**Authors:** Alejandro Garcia Garcia, Sujeethkumar Prithiviraj, Deepak Bushan Raina, Tobias Schmidt, Sara Gonzalez Anton, Laura Rabanal Cajal, David Hidalgo Gil, Robin Kahn, Magnus Tägil, Paul Bourgine

## Abstract

Tissue engineering strategies predominantly consist of the autologous generation of living substitutes capable of restoring damaged body parts. Persisting challenges with patient-specific approaches include inconsistent performance, high costs and delayed graft availability. Towards developing a one-for-all solution, a more attractive paradigm lies in the exploitation of dedicated cell lines for the fabrication of human tissue grafts. Following decellularization, this new class of biomaterials relies on the sole extracellular matrix and embedded growth factors instructing endogenous repair. This conceptual approach was previously validated using a custom mesenchymal line for the manufacturing of human cartilage, exhibiting remarkable osteoinductive capacity following lyophilization. Key missing criteria to envision clinical translation include proper decellularization as well as stringent assessment of both immunogenicity and regenerative performance. Here, we report the engineering and subsequent decellularization of human cartilage tissue with minimal matrix impairment. Ectopic evaluation in immunocompetent and immunocompromised animals reveal preservation of osteoinductivity predicted by macrophage kinetic of polarization. By establishing in vitro human allogeneic co-culture models, we evidenced the immuno-evasive properties of cell-free human cartilages, controlling macrophages and dendritic cells maturation as well as T cell activation. Lastly, regenerative performance was stringently assessed in an immunocompetent rat orthotopic model whereby decellularized human cartilage grafts achieved morphological and mechanical restoration of all critical-sized femoral defects. Taken together, our study compiles robust safety and efficacy pre-requisites prompting a first-in-human trial for engineered and decellularized human tissue grafts.

## Introduction

Musculoskeletal traumas remain the first cause of disability affecting over 1.7 billion people worldwide(1). Bone injuries and disorders account for a large fraction of this burden(2), creating a substantial demand for effective treatments that an aging population only render more acute. Tissue engineering has led to the development of a myriad of biomaterials and bioengineered substitutes shown to stimulate bone regeneration(3). To enhance regenerative performance, a significant breakthrough involved incorporating developmental engineering principles into design strategies. This approach harnesses the natural process of endochondral ossification, where cartilage tissue is engineered to act as a transient template for bone formation(4). To this end, Hypertrophic cartilage (HyC) can be generated *in vitro* by differentiating various cell sources, such as primary human mesenchymal stem/stromal cells (hMSCs) on scaffolding materials(5). Upon implantation, the engineered HyC is gradually remodeled into mature bone matrix through the differentiation of chondrocytes into osteoblasts and the recruitment of endogenous osteoprogenitor cells(6). Despite showing great promise in terms of graft integration and bone tissue restoration, key challenges inherent to autologous approaches remain to be overcome. These include donor site morbidity, limited availability and variability, as well as inconsistencies in protocols for differentiating hMSCs into chondrocytes(7, 8). These factors add layers of complexity, making it difficult to standardize and optimize the process for consistent and effective clinical outcomes.

Limits of living strategies steered the development of an acellular alternative, whereby the regenerative process would solely rely on the graft extracellular matrix and embedded factors. Our group has made significant contributions to this conceptual approach by engineering cartilage tissues *in vitro* using the MSOD-B line as a source of immortalized hMSCs(9). We demonstrated the scalable production of human cartilage that retains strong osteoinductive properties even after lyophilization. This work established the foundation for manufacturing human tissue substitutes as off-the-shelf biomaterials capable of guiding and promoting bone repair.

While providing a strong proof-of-concept, the journey to clinical translation remains challenging due to the lack of complete decellularization and stringent performance assessments. The pre-clinical validation of cell-based but cell-free grafts is further complicated by the fact that this new class of biomaterials is yet to be fully defined by regulatory agencies(1, 10). Additionally, the suitability of standard pre-clinical models remains to be determined, as often falling short on replicating the complexity of the human immune system and physiological responses(11, 12).

In this study, we aim at achieving the generation of decellularized HyC and at determining both its immunogenicity and bone-repair capacity. We first propose to identify an effective decellularization protocol preserving the engineered tissue, followed by the evaluation of the resulting osteoinductive properties and immune recruitment using immunodeficient and immunocompetent mouse models. We further design *in vitro* human immunogenicity assays, in which allogeneic antigen presenting cells are analyzed for their ability to activate an immune response following exposure to HyC. Finally, grafts will be tested as a callus replacement biomaterial in a critical-sized femoral defect model. By providing both safety and efficacy data, our work aims at prompting the clinical translation of cell-line derived human tissue as grafts engineered to instruct endogenous repair.

## Results

### Cell-line engineered human cartilage can be efficiently decellularized with minimal matrix impairment

To identify an optimal decellularization protocol offering both efficient DNA removal and preservation of tissue integrity, 12 protocols were evaluated including distinct detergent treatments (Triton X-100, SDC, and SDS), hypertonic and hypotonic baths and DNase digestion time (**Supplementary Figure 1A**). Human hypertrophic cartilage pellets were in vitro engineered as previously described(9) using the MSOD-B line (**Figure 1A**), an immortalized hMSCs capable of chondrogenesis. After lyophilization, tissues were exposed to various decellularization protocols and readouts consisted in measurements of total DNA and total glycosaminoglycans (GAGs) amount for decellularization efficiency and matrix preservation respectively. Three protocols (Protocol 6, 9 and 12) were selected based on their reduced impact on GAGs content and efficient DNA removal and further evaluated by assessing BMP-2 and total collagen preservations (**Supplementary Figure 1B**), as well as histological analysis (**Supplementary Figure 1C**) of the resulting cartilage pellets. This allowed the identification of one protocol demonstrating efficient decellularization with good preservation of graft constituents (Protocol 12, **Supplementary Figure 1A**). The resulting decellularized hypertrophic cartilage (D-Hyc) were further analyzed and directly compared to its non-decellularized but lyophilized counterpart (L-HyC). Scanning electron microscopy radiographs revealed an increase surface porosity and exposure of the fibrillar matrix existing within the D-HyC tissue, with no cells visible on the surface (**Figure 1B**). Taking advantage of the MSOD-B cells GFP expression (**Figure 1A**), confocal imaging of cartilage tissues revealed a homogenous cellular distribution in L-HyC. Instead, D-HyC staining evidenced an absence of both GFP and DAPI (nuclei) signals, confirming the decellularization efficacy. Quantification of both DNA and GFP levels in digested tissues corroborated the successful decellularization, with D-HyC samples exhibiting negligible DNA levels (0.039µg/mg of dry weight) as compared to L-HyC (13.4 µg/mg of dry weight). This corresponded to a >99.7% decellularization efficiency (**Figure1C**), a residual DNA traces were below the suggested threshold defined in the field(13). To quantify the impact on the cartilage ECM, we further assessed histologically the presence of GAGs, collagen type-2 and collagen type-X (**Figure 1D**). While both L-HyC and D-HyC display features of hypertrophic cartilage matrix, the D-HyC samples showed a noticeable GAGs reduction and a disrupted collagen-2/collagen-X networks as compared to L-HyC. This was further confirmed by a quantitative Blyscan assay, revealing a 17.6% loss in GAG content compared to non-decellularized controls (**Figure 1E**). In contrast, analysis of other structural proteins indicated the preservation of total collagens composition **(Figure 1E)**. Similarly, the levels of BMP-2 embedded within the matrix remained unaffected (**Figure 1E)**.

**Figure 1:**
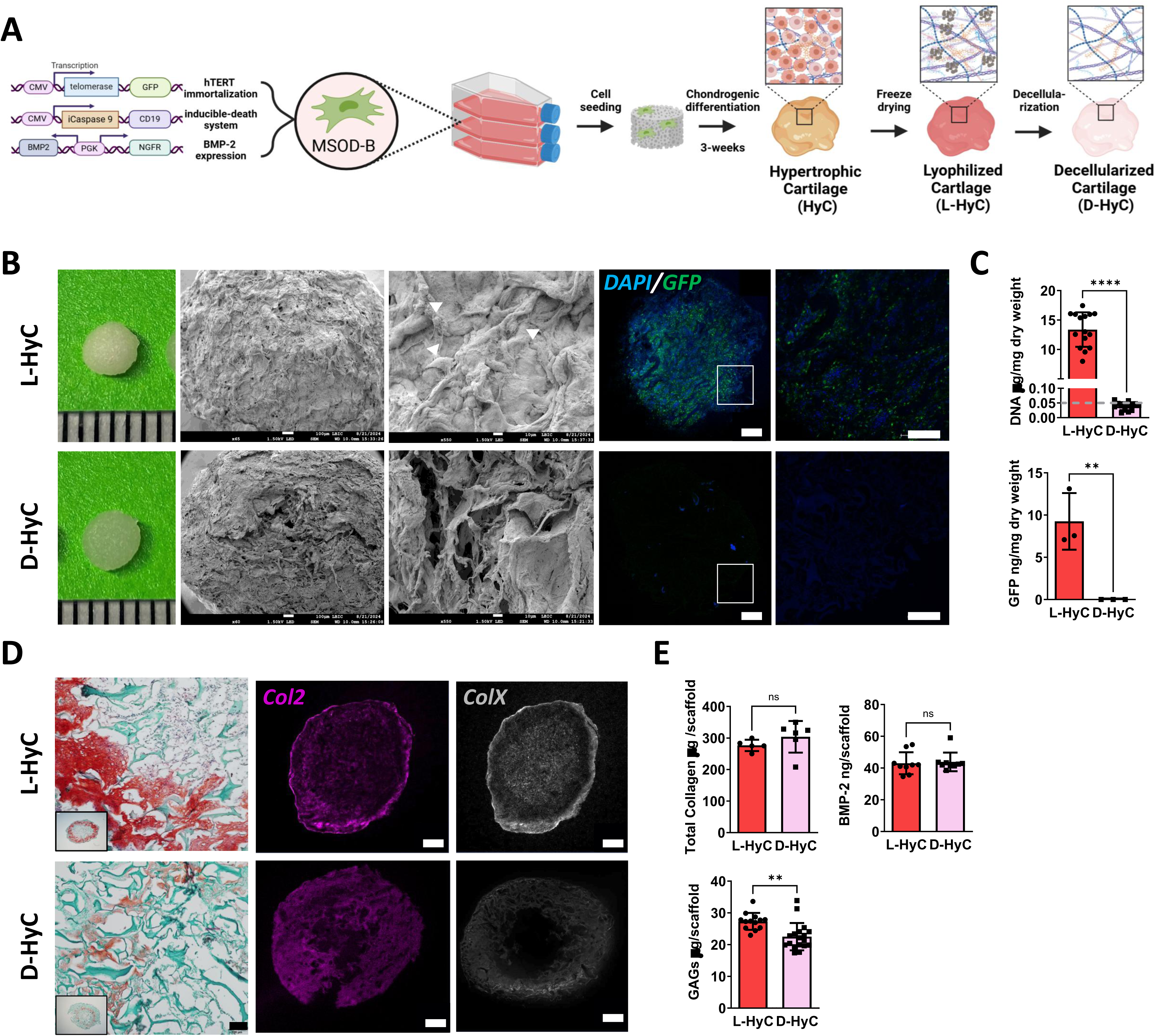
Cell-line engineered human cartilage can be efficiently decellularized with minimal matrix impairment. **A.** Experimental scheme for the generation of decellularized human cartilage grafts. **B.** From left to right: Macroscopic representation; scanning electron microscopy pictures of graft surfaces (arrows indicate cells), Scale bar = 1mm and 100μm; and immunofluorescence confocal images of grafts stained for DAPI and GFP. **C.** From top to down quantification of total DNA (n=15) and GFP (n=3) remnants per mg dry weight of pellets. Dashed line represent the reported threshold for efficient decellularization (0.05 μgDNA/mg dry weight) established in the literature **D.** From left to right: Safranin-O staining and immunofluorescence images of human grafts stained for Col2 and ColX, Scale bar = 1mm. **E.** Quantification of GAGs (n=15), total collagen (n=6) and BMP-2 content (n=9) on decellularized and lyophilized grafts. Graphs represent mean ± standard deviation (SD), \*\**p* ≤ 0.01, \*\*\**p* ≤ 0.001, determined by two-tailed unpaired *t*-test.

Taken together, we here report the identification of an efficient decellularization protocol, affecting GAGs content but preserving collagens and BMP-2 compositions.

### Decellularized human cartilage grafts fully remodeled into bone organs by ectopic priming of endochondral ossification in immunodeficient mice

To assess whether the decellularization process affects the capacity of human-engineered cartilage grafts to prime endochondral ossification, we implanted lyophilized (L-HyC) and decellularized (D-HyC) grafts ectopically into immunodeficient mice (**Figure 2A**). Early time-point analysis revealed similar remodeling of L-HyC and D-HyC, with progressive cell colonization. By day 10, the cartilage matrix has been fully degraded by host cells (Safranin-O) accompanied by intensification of collagen matrix deposition, as confirmed by Masson’s trichrome staining (MT) (**Figure 2B**).

**Figure 2:**
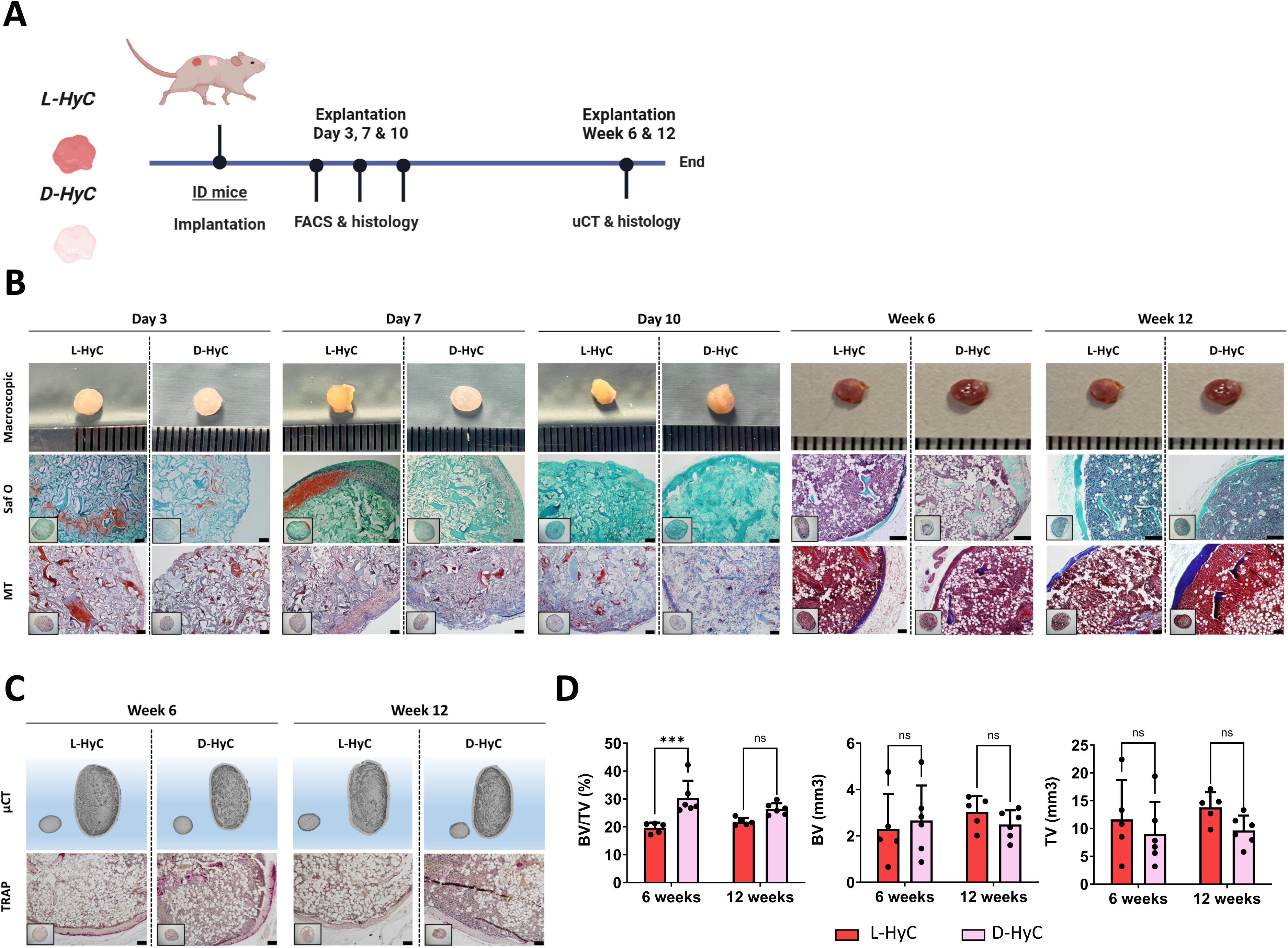
Decellularized human cartilage grafts fully remodeled into bone organs by ectopic priming of endochondral ossification in immunodeficient mice. **A.** Experimental scheme for the *in vivo* ectopic osteogenic assessment of L-HyC and D-HyC grafts. **B.** From top to bottom, representative macroscopic images (scale bar=1mm), histological images of Safranin-O and Masson Trichrome staining of explanted tissues (Scale bar =100µm). **C.** From top to bottom, representative µCT 3D reconstruction and TRAP staining of explanted tissues at 6- and 12-weeks (Scale bar =400µm). **D.** BV/TV (%), BV (mm^3^) and TV (mm^3^) of µCT performed over explanted bones at 6- and 12-week post-implantation (n≥5). The graphs represent mean ± standard deviation (SD), *** *p* ≤ 0.001, determined by Two-way ANOVA.

Remarkably, by 6-weeks post-implantation both L-HyC and D-HyC grafts fully remodelled into mature bone and bone marrow tissues (**Figure 2B**). Safranin-O and Masson’s trichrome stainings demonstrated the replacement of cartilage with mature bone tissue, including marrow infiltration and the formation of a cortical bone layer with trabecular structures (**Figure 2B**). Micro-CT analyses confirmed macroscopical observations while TRAP staining revealed active osteoclast-mediated remodelling at 6-weeks, consistent with mature bone turnover (**Figure 2C**). At 12-weeks post-implantation, tissues persist in vivo marking a definite remodeling with no detectable osteoclastic activity. Micro-CT radiographs allowed quantification of bone mineralization (**Figure 2D**) and tissue volume at 6- and 12-weeks post-implantation. This confirmed the similar performance of both L-HyC and D-HyC tissues, with only a superior bone volume / tissue volume ratio (BV/TV) in D-HyC at the 6-week timepoint. By 12-weeks, no significant differences in both bone and tissue volume were found between groups.

These results demonstrate the strong osteoinductive properties of engineered human cartilage grafts and evidences the intrinsic bone forming capacity of fully decellularized human cartilage tissue.

### Decellularized human cartilage does not induce robust ectopic bone formation in immunocompetent mice

Given the efficient decellularization achieved, we aimed to evaluate whether our grafts could similarly achieve ectopic bone formation when exposed to a fully functional immune system. To this end, L-HyC and D-HyC tissues were implanted into C57BL/6 and tissue remodeling was assessed across various time-points (**Supplementary Figure 2A**). Within the first 10-days post-implantation, the cartilage matrix is progressively degraded in both L-HyC and D-HyC (**Supplementary Figure 2B**). Concomitantly, a thick collagen layer was observable at the periphery of the implants, suggesting an encapsulation.

At later time points (6- and 12-weeks), none of the retrieved samples exhibited frank bone formation histologically (**Supplementary Figure 2B**), mostly consisting of fibrous collagen matrix. While D-HyC and L-HyC exhibited similar histological features, their retrieval rates largely differed. Most L-HyC samples were fully degraded by the host, with only 40% and 20% of tissues being recovered at 6- and 12-weeks respectively. In turn, 80% of D-HyC were found at both 6- and 12-weeks post-implantation. Among retrieved samples, TRAP stainings fail at detecting any bone remodeling but small areas of mineralization could be identified in D-HyC samples only (**Supplementary Figure 2C** and 2D).

These findings highlight the immunogenic nature of the human matrix components within engineered cartilage grafts, leading to rejection at ectopic site in immunocompetent mice. The removal of human cells and DNA suggest a reduction of the immune reaction associated with higher persistence of D-HyC tissues, although failing at priming endochondral ossification.

### Immune prints of engineered human cartilage correlate early M0 to M2 polarization with successful ectopic bone formation

Given the successful bone formation observed in immunodeficient (ID) and failure in immunocompetent (IC) animals, we seek to further investigate the underlying inflammatory properties of L-HyC and D-HyC grafts in both ID and IC mice. To do so, we performed a temporal analysis of immune recruitment in the first days (3, 7 and 10) post-implantation, corresponding to the reported acute inflammatory phase preceding remodeling(14, 15).

The total number of cells recruited to implants was shown to gradually increase overtime (**Supplementary Figure 3A**), oscillating between 0.5 to 1M cells per graft in both ID and IC settings. In ID animals, a consistent superior recruitment was observed in L-HyC as compared to D-HyC. A similar trend was observed in IC with L-HyC recruiting more cells, although not reaching significance. This difference in cell recruitment largely accounted for blood/immune cell infiltration (CD45+, **Figure 3A**), particularly marked at days 7 and 10 post-implantation. To further identify and discriminate the different recruited immune populations, a comprehensive panel of phenotypic markers was set for flow cytometry analysis (**Supplementary Figure 3B**). This allowed compiling “immune prints”, capturing the immune composition of implanted grafts as a snapshot in respective timepoints and settings (**Figure 3B, Supplementary Figure 3C**). These radar plots consistently identified macrophages (MΦ), natural killer cells (NKs), dendritic cells (DCs), T cells (Tcs), and B cells (Bcs) within the implanted tissues. Strikingly, the immune prints of L-HyC and D-HyC revealed to be very similar across all timepoints as well as between ID and IC animals. Macrophages were identified as the most abundant recruited cell type, with their frequency peaking on day 10 post-implantation. While both grafts immune prints closely aligned, D-HyC exhibited a statistically lower macrophages frequency at day 3 and increased NKs at day 10 in ID animals. In IC, D-HyC exhibited statistically higher DCs at day 3, higher T-cell frequency at day 7 and day 10 and increased NK at day 10. In the other hand, they exhibited lower frequency of macrophages at day 10. (**Supplementary Figure 3C**).

**Figure 3:**
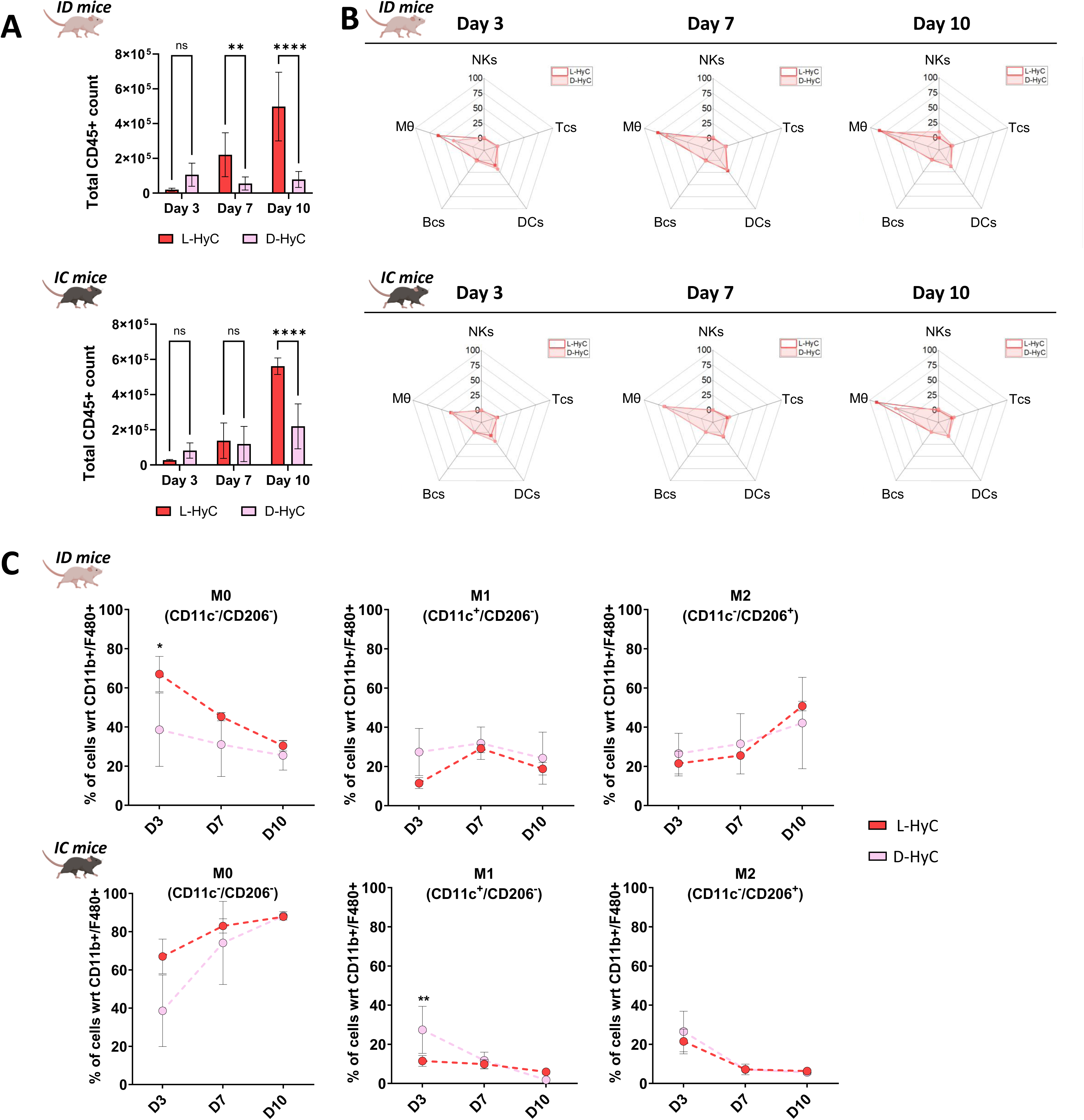
Immune prints of engineered human cartilage correlate early M0 to M2 polarization with successful ectopic bone formation. **A.** Total cell number captured in explanted tissues after 3-, 7-, and 10-days post-implantation in immunodeficient (Top) and immunocompetent (Bottom) animals. The graphs represent mean ± standard deviation (SD), \*\**p* ≤ 0.01, \*\*\*\**p* ≤ 0.0001, determined by Two-way ANOVA. **B.** From right to left, representative radar plots representing percentages of early time recruitment of B-cells, T-cells, Dendritic cells, Natural Killers and Macrophages captured in explanted tissues after 3-, 7-, and 10-days respectively in immunodeficient (Top) and immunocompetent (Bottom) animals. **C.** Percentage of CD11b^+^ and F4-80 ^+^ macrophage subtypes in L-HyC and D-HyC grafts explanted at 3-, 7- and 10-days post-implantation analyzed by flow cytometry in immunodeficient (Top) and immunocompetent (Bottom) animals. Error bars: The graphs represent mean ± standard deviation (SD), \**p* ≤ 0.1, \*\*\**p* ≤ 0.001, determined by Two-way ANOVA. N≥3 independent experiments, 3 pooled samples per animal (n ≥9).

Based on the observed predominant presence of macrophages, we further explored their distinct polarization states (**Figure 3C**). Importantly, macrophage polarization dynamics were similar between L-HyC and D-HyC grafts. However, substantial pattern differences could be observed by comparing ID and IC animals. In ID, we observed an initial dominant unpolarized (M0) state which progressively declined overtime. Concomitantly, a progressive increase in pro-regenerative macrophages (M2) was observed reaching its peak by day 10. Instead, the pro-inflammatory M1 population remain stable, suggesting a M0 to M2 polarization process.

Instead, in IC animals we report a progressive increase in M0 macrophages, reaching over 88% of total macrophages by Day 10 (**Figure 3C**). In sharp contrast with the ID animals, no increase in M1 nor M2 populations was observed, those populations even declining over the 10 days observational course.

Taken together, the early immune profiling of L-HyC and D-HyC exhibit comparable inflammation which aligns with the similar osteoinductive performance in ID and IC. By further correlating immune profile and bone formation, our data suggests that a M0-to-M2 polarization is required and predictive of successful ectopic ossification, as observed for both L-HyC and D-HyC. The L-HyC and D-HyC immune prints suggest a similar immunogenicity of our grafts despite the decellularization. Alternatively, it may evidence limits of *in vivo* models in assessing immune reaction caused by human grafts.

### Human allogeneic in vitro assays reveal minimal macrophage polarization and absence of T cell activation by decellularized human cartilage grafts

Towards mimicking an inflammation scenario upon clinical implantation, we investigated the immunogenicity of engineered cartilage grafts in a fully human setting by design of in vitro assays. To this end, we first developed a setup whereby monocytes from healthy individuals (n = 6 donors) were isolated and differentiated towards macrophages (**Figure 4A**). Their polarization status was assessed post-differentiation, when exposed to L-HyC or D-HyC. As positive polarization controls, macrophages were exposed to LPS and IFN-γ to form M1 macrophages, or IL-10 and dexamethasone for M2 macrophages induction. To maximize cell exposure to the graft material, both L-HyC and D-HyC were cryo-milled into a fine powder and subsequently added to the culture medium. After macrophage differentiation, flow cytometry revealed an increase in pro-inflammatory M1 markers (CD80 and CD86) upon exposure to L-HyC (**Figure 4B**). Instead, D-HyC did not induce a significant M1 polarization, as compared to macrophages not exposed to any HyC materials (Untreated). Moreover, none of the grafts were shown to induce a M2 polarization (CD163 and CD206, **Figure 4B**). Comparing polarization patterns across donors (**Supplementary Figures 4A** and 4B), we observed variable M1 activation in the L-HyC group, whereas D-HyC consistently led to lower activation of pro-inflammatory macrophage markers. Similarly, while L-HyC generally induced a stronger M2 response compared to D-HyC, this effect was highly donor-dependent.

**Figure 4:**
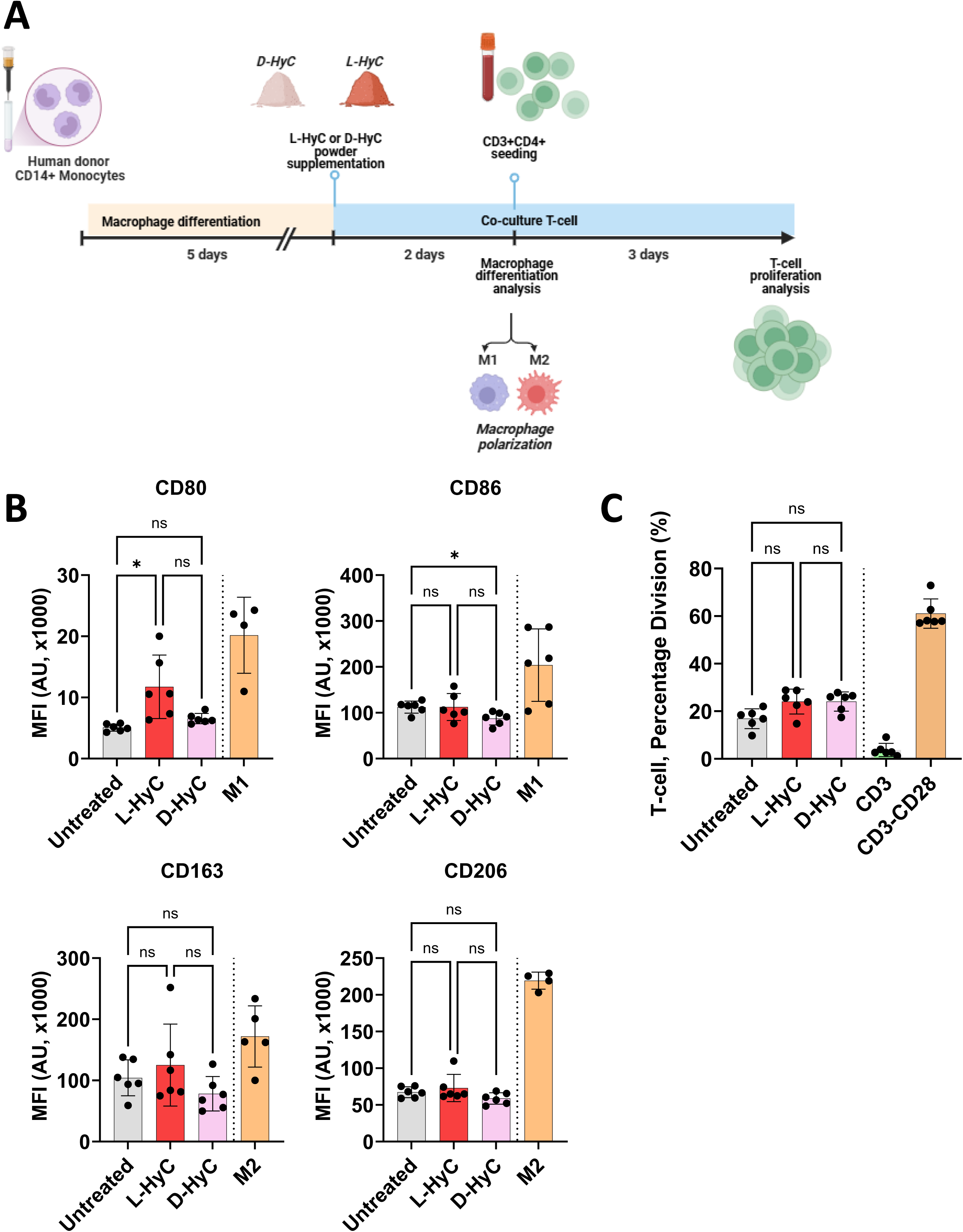
Human allogeneic in vitro assays reveal minimal macrophage polarization and absence of T cell activation by decellularized human cartilage grafts. **A.** Experimental scheme of the direct effect of powdered L-HyC and D-HyC on macrophage polarization and indirect T-cell activation potential. **B.** From top to bottom, total CD80+ and CD86+ MFI indicating M1 polarization; and total CD163+ and CD206+ MFI indicating M2 polarization, after 5-days co-culture with either L-HyC or D-HyC powdered cartilages. The graphs represent mean ± standard deviation (SD), \**p* ≤ 0.1 determined by repeated measures one-way ANOVA(6 donors, n=3 per donor). **C.** CD3+ percentage division assessed by FACS after co-culture of 5x10^4^ CD3+ cells with 2.5x10^3^ macrophages co-cultured with either L-HyC or D-HyC powdered cartilages . The graphs represent mean ± standard deviation (SD), determined repeated measures one-way ANOVA, statistical significance set at *p < 0,05.*(6 donors, n=3 per donor).

Next, we evaluated the capacity of resulting macrophages to differently induce T-cell activation/proliferation after exposure to L-HyC or D-HyC (**Figure 4C**). As a positive control, T-cells were treated with anti-CD3 and anti-CD28, where CD28 acts as co-stimulatory signal leading to full T-cell receptor (TCR) activation. Anti-CD3 only was used to directly stimulate TCR, leading to incomplete activation. After three days, only the CD3-CD28 group exhibited substantial T-cell proliferation (60%). In contrast, T-cells co-cultured with macrophages exposed to either L-HyC or D-HyC showed minimal activation (**Figure 4C**). These results indicate that macrophages exposed to L-HyC or D-HyC, despite and increase of co-stimulatory molecules following L-HyC stimulation, do not significantly induce T-cell proliferation. It further reveals that DNA and cellular remnants in L-HyC did not cause an increased inflammatory macrophage-driven response. Furthermore, the absence of significant differences between untreated and construct-exposed groups suggests that neither L-HyC nor D-HyC per se are capable of triggering T-cell activation. In fact, co-culturing T-cells with M1 or M2 macrophages (**Supplementary Figure 4C**) resulted in the same level of T-cell activation as observed with L-HyC and D-HyC tissues. This suggests that the engineered grafts do not further amplify macrophage-driven inflammation beyond the baseline macrophage activation state. Instead, T-cell activation is primarily driven by CD3 signaling rather than macrophage phenotype and associated inflammatory cytokines. In fact, our findings indicate that HyC matrices do not push macrophages toward a highly inflammatory M1-like state nor enhance their antigen presentation capacity.

Overall, our data revealed that the decellularization of human cartilage grafts reduced M1 polarization while minimizing donor-to-donor variability. Unlike findings from mouse studies, no M2 polarization was observed in our *in vitro* assays. Notably, engineered L-HyC and D-HyC materials minimized T-cell activation induced by macrophages.

### Human engineered cartilage grafts do not elicit a human adaptive immune response in vitro while exhibiting immunosuppressive properties

In light of the observed macrophages response, we further assessed whether an adaptive immune response could be mounted after exposure to L-HyC or D-HyC. We thus isolated full mononuclear cells (PBMCs) -containing T cells- and similarly exposed them to grinded human cartilages for 5 days (**Figure 5A**). Phytohemagglutinin (PHA) was used as a positive control ensuring T cell proliferation, whereas PBMCs not exposed to any human cartilage was set as a control group (Untreated).

**Figure 5:**
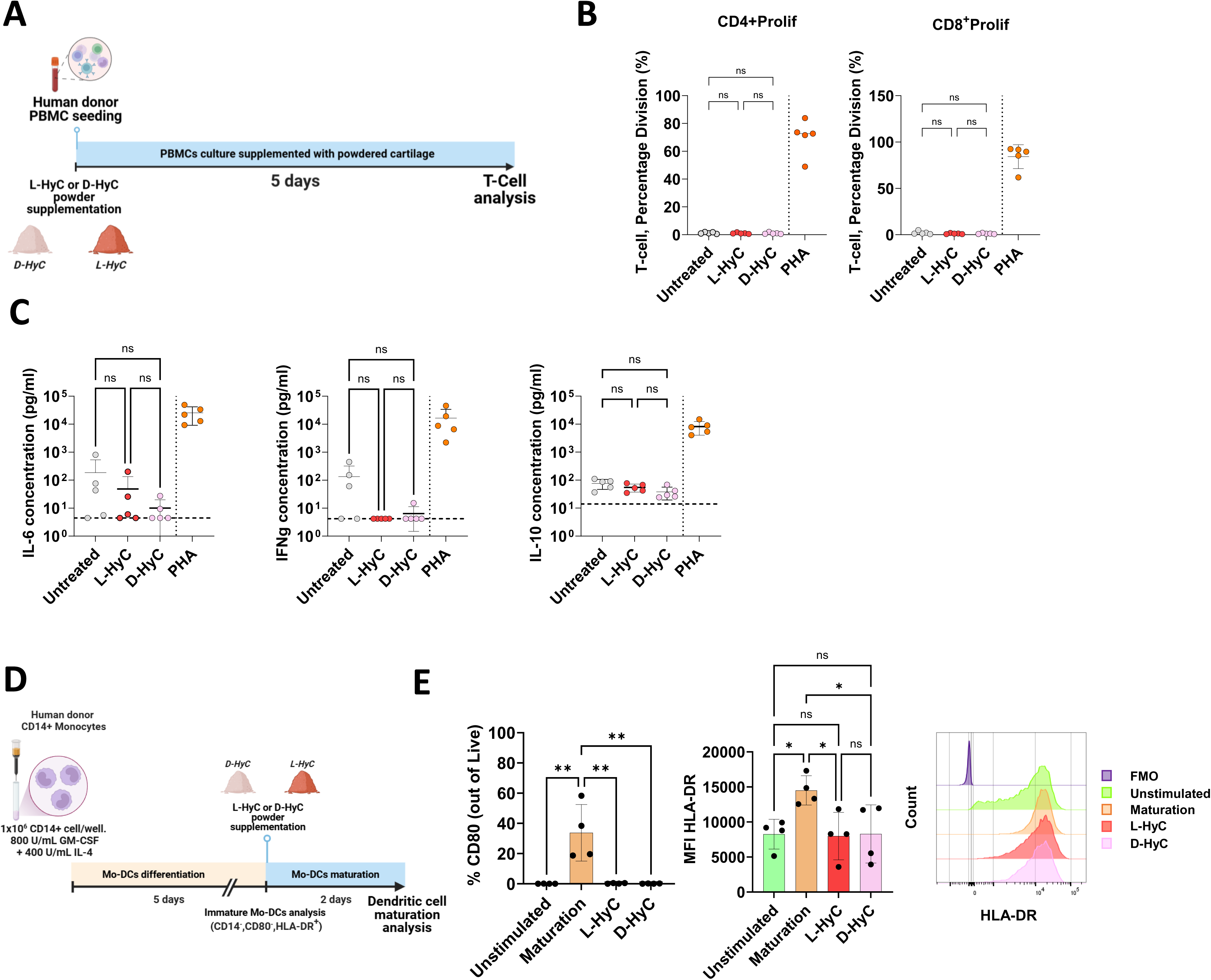
Human engineered cartilage grafts do not elicit a human adaptive immune response in vitro while exhibiting immunosuppressive properties. **A.** Experimental scheme of the direct effect of powdered L-HyC and D-HyC on T-cell activation when directly co-cultured with PBMCs for 5 days **B.** From right to left, percentage of activated CD3+CD4+ and CD3+CD8+ T-cells. The graphs represent mean ± standard deviation (SD), determined by repeated measures one-way ANOVA, statistical significance set at *p < 0,05.* C. Pro-inflammatory cytokines released by PBMCs after 5 days co-culture with either L-HyC or D-HyC powdered cartilages. The graphs represent mean ± standard deviation (SD), determined by repeated measures one-way ANOVA, statistical significance set at *p < 0,05*. The horizontal dashed-lines indicates the detection threshold of the ELISA assay. **D.** Experimental scheme of the assessment of dendritic cell maturation when cultured with either L-HyC or D-HyC powdered cartilages or maturation media in combination with powdered cartilages for 48H **E.** From left to right, percentage of CD80+ and total HLA-DR+ MFI of dendritic cells dendritic cells when exposed with either L-HyC, D-HyC powdered cartilages or maturation media in combination with powdered cartilages for 48H. The graphs represent mean ± standard deviation (SD), \*\**p* ≤ 0.01, determined by Ordinary one-way ANOVA.

As anticipated, the PHA condition strongly induces T cell proliferation with 74% and 87% of CD4 or CD8 T-cells respectively entering division (**Figure 5B**). In stark contrast, PBMCs exposed to the human cartilage epitopes were unable to activate T cells (**Figure 5B**). Strikingly, both CD4+ and CD8+ did not enter proliferation, with a percentage of T cell division similar to the untreated group (**Figure 5B**). Inflammatory cytokines (IFN-γ, IL-6 and IL-10) measured in the co-culture medium confirmed those observations (**Figure 5C**), with L-HyC and D-HyC values equivalent to those from the Untreated group. Taken together, these data confirmed the low immunogenicity induced by L-HyC and D-HyC in vitro (**Figure 5C**).

Using flow cytometry, we further analyzed T-cell activation status by assessing subtype distribution based on surface marker expression. As anticipated, the PHA group exhibited the largest activation of various T cell subsets (**Supplementary Figure 5A**). However, no differences across HyC groups could be observed, with a detected but limited percentage of CD4⁺CD25⁺, CD4⁺HLA-DR⁺, CD8⁺CD25⁺ and CD8⁺HLA-DR⁺ T-cell subsets (**Supplementary Figure 5A**). This suggests an immuno-evasive effects of HyC regardless of their decellularization status.

The absence of T cell activation questioned whether the human cartilage grafts can impair the function of antigen presenting cells. As DCs accounts for professional antigen presenting entity, we assessed whether their maturation could be affected by the L-HyC or D-HyC matrices. To this end, monocytes were first isolated from healthy donors and pre-differentiated into immature DCs (**Figure 5D**). At that stage, over 99% of the cells expressed typical DCs markers (CD14^-^HLA-DR^+^) but not activated ones (<1% of CD80+) (**Supplementary Figure 5B**). Immature DCs were then exposed to a maturation media (positive control) or to the L-HyC and D-HyC grinded proteins to assess their capacity to induce DCs maturation. The maturation media led to a robust DCs activation with 33% of cells expressing the CD80 markers, in conjunction with an increased in mean HLA-DR expression intensity. (**Figure 5E**). In contrast, when exposed to L-HyC or D-HyC, immature DCs fail at further differentiating, with no detectable CD80 expression and no increased in HLA-DR intensity (**Figure 5E**).

In summary, those findings evidenced the low immunogenic nature of cell-free human cartilage grafts. Using *in vitro* human-based assays, the decellularization process did not significantly reduce the inflammatory status as L-HyC and D-HyC presented similar-to-identical-properties. Interestingly, despite the absence of living cells, the cartilage ECMs displayed immunoregulatory features linked with reduced T cell activation by PBMCs and impairment of DC maturation.

### Decellularized hypertrophic cartilage promote full healing of critical-sized femoral defect in immunocompetent rats

After demonstrating potent osteoinductivity and reduced immunogenicity, we last assessed the repair capacity of our human engineered graft in a pre-clinically relevant setting. To this end, D-HyC tissues were implanted into a critical-sized femoral defect in an immunocompetent rat model (**Figure 6A**). To fill the 5mm defect, 3-pellets were implanted at the surgical site (**Figure 6B**), with the graft diameter approximating the diameter of the femur. As negative control, an empty defect was created. The repair capacity was monitored for up to 12 weeks by micro-computed tomography (micro-CT) followed by histological assessments.

**Figure 6:**
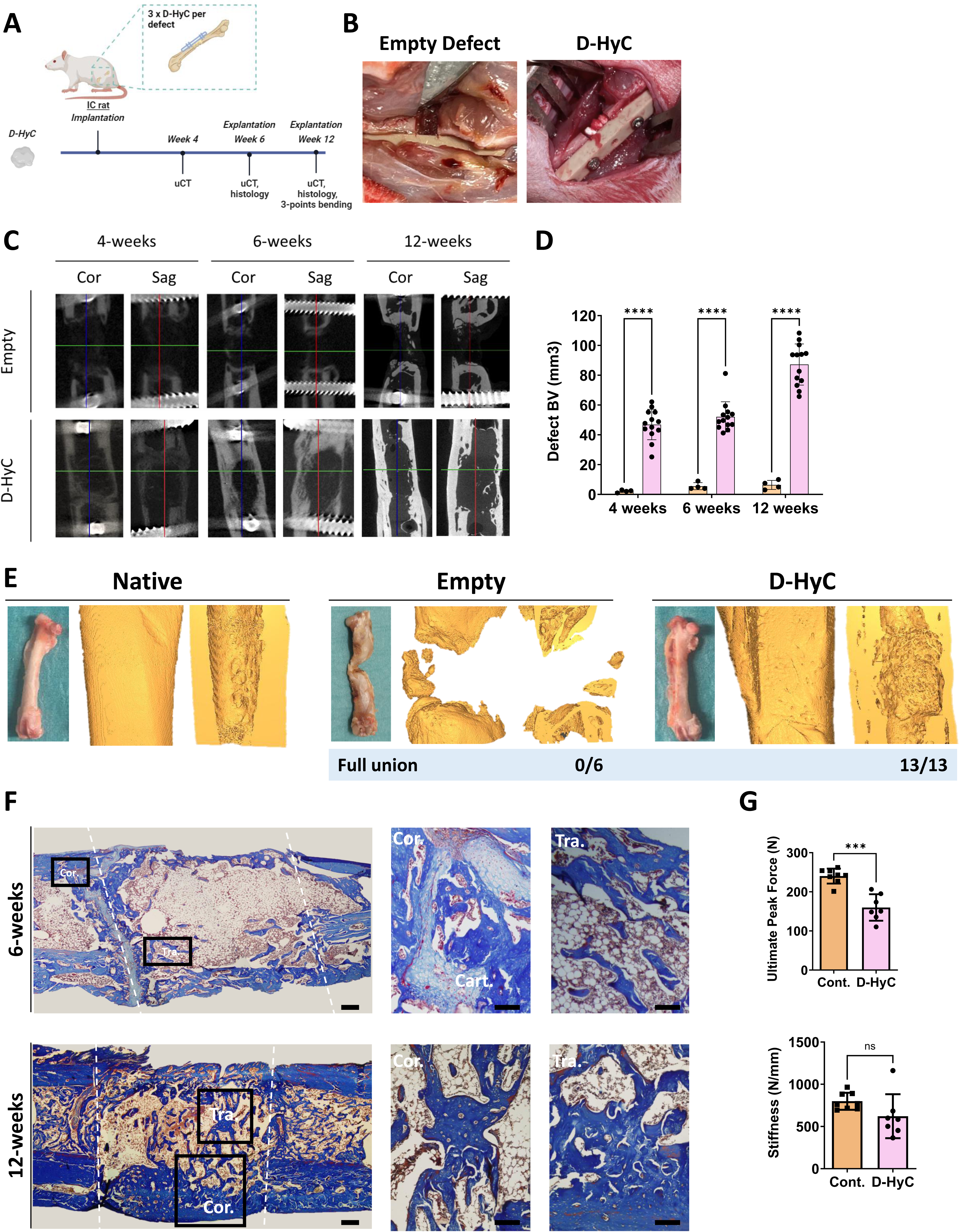
Decellularized hypertrophic cartilage promote full healing of critical-sized femoral defect in immunocompetent rats. **A.** Experimental scheme of the assessment of bone regenerative potential of decellularized tissue engineered cartilages assessed in a critical full-sized femoral defect in immunocompetent rat. **B.** From left to right, macroscopic images of the empty critical full-sized defect and the defect fulfilled with a total of three engineered hypertrophic cartilage tissues. **C.** Micro-CT based representative 2D coronal (left) and sagittal (right) views from the middle of the defect after 4-, 6- and 12-weeks for empty groups and regenerated femurs implanted with D-HyC. **D.** Quantification of micro-CT based bone volume (BV) in the defect (region of interest height = 4.5 mm) for empty groups and regenerated femurs implanted with D-HyC after 4-,6- and 12-weeks of implantation. The graphs represent mean ± standard deviation (SD), \*\*\*\**p* ≤ 0.0001, determined by Two-way ANOVA. **E.** From left to right, representative macroscopic image of an explanted regenerated femur at 12-weeks; representative 3D images of the entire defect area in both whole and cross-sectional view. **F.** From left to right, representative histological images of Masson Trichrome staining of explanted femurs completely regenerated after 12-weeks implantation with D-HyC; and a magnification of a region of interest of a cortical (cor.) and a trabecular (tra.) bone structure, Scale bar = 500 and 100 µm respectively. **G.** From left to right, Ultimate Peak Force (N) and total Stiffness (N/mm) for empty groups and regenerated femurs implanted with D-HyC after 12-weeks of implantation (n=7). The graphs represent mean ± standard deviation (SD), \*\*\**p* ≤ 0.001, determined by Two-way ANOVA.

As early as 4 weeks, micro -CT revealed significant mineralization in D-HyC implants as compared to empty controls (**Figure 6C**). Quantification across timepoints confirmed the progressive repair, with potential defect bridging visualized at 6 weeks in 12 out of 13 animals (**Figure 6C** and **6D**). Bone/mineralized volume in the defect region constantly increased reaching 87mm^3^ by 12 weeks. Femurs from animals were then explanted at 6 and 12 weeks for histological analysis. Remarkably, upon explantation, macroscopic and micro-CT evaluation revealed full-bridging in the D-HyC treated group (**Figure E**). This was the case for all operated animals (13/13), with D-HyC femurs resembling the non-operated contralateral ones (**Supplementary Figure 6**). Instead, empty controls did not exhibit robust repair (**Figure 6E**). 3D reconstructions and cross-sectional views confirmed robust bone regeneration in the D-HyC group only and bridging of the defects (**Figure 6E**). To confirm frank de novo bone formation, we performed histological analysis using Masson Trichrome staining. At 6-weeks post-implantation, an extensive bone formation was observed in the defect area, bridging with the adjacent cortical structures (**Figure 6F**). At that timepoints, a cartilage layer could be observed at the native bone/defect interface, indicating an ongoing endochondral ossification process. Trabecular structures and restoration of the bone marrow cavity were also evidenced at that timepoint. At 12-week, a more extensive bone trabecular network was observed, along with a thickening of the cortical structure (**Figure 6F**). At that stage, no cartilage remnants were observed, suggesting a definite regeneration. We further compared the bridging performance of D-HyC with published grafting strategies evaluated in the same immunocompetent rat femoral model (**Supplementary Figure 7**)(16–18). Only a high dose of BMP2 and syngeneic transplantation (rat living cartilage graft) match the performance of D-HyC at 12 weeks post-implantation, with consistent defect bridging.

Finally, we performed a biomechanical evaluation of retrieved femurs, towards assessing the extent of their functional restoration. D-HyC treated femurs 12-week post-implantation were tested, while non-operated contralateral ones served as positive controls. Using the ultimate peak force assay, D-HyC femurs were shown to achieve 67% of the contralateral femur resistance to fracture (160N versus 240N, for D-HyC versus control respectively, **Figure 6G**). Instead, stiffness measurements indicated no statistical differences between D-HyC and control femurs.

## Discussion

In this study, we report the successful generation of decellularized HyC, retaining remarkable osteoinductive properties predicted by early M0 to M2 polarization. Independently from the decellularization process, HyC demonstrated potent immunoregulatory properties, characterized by minimal human allogeneic T cell activation and reduced maturation. When implanted in a rat critically sized defect, D-HyC led to full femoral regeneration with morphological and mechanical bone restoration.

The decellularization of engineered tissues present the attractive opportunity to create off-the-shelf grafting solutions, eliminating the need for immuno-matching(19, 20). To achieve an effective removal of immunogenic cell debris, existing protocols often compromise the composition and thus the performance of engineered tissues. This challenge is particularly pronounced in cartilage due to its dense cartilage matrix(21). Physical methods often disrupt its cellular membranes and nuclei, but it may not completely remove all cell components, while chemical processing can disrupt collagen network and cause significant loss of GAGs and growth factors. A tailored combination of chemical and physical methods is needed for effectively decellularizing such tissues(22). In this study, we successfully established a decellularization protocol using an optimized combination of detergents (SDS), hypertonic/hypotonic buffers, and DNase treatment. SDS is widely used in cartilage decellularization at concentrations ranging from 1–2% for 24–72 hours(23, 24). However, due to its cytotoxic effects, which can compromise the biocompatibility of decellularized grafts(25), we optimized the protocol by reducing SDS concentration to 0.5% and limiting exposure to 12 hours. To further improve efficiency while preserving ECM integrity, we implemented a stepwise approach, sequentially applying treatments rather than exposing the tissue to all reagents simultaneously. While hypertonic and hypotonic baths are typically used to lyse cells and precipitate cellular debris, we additionally utilized the hypertonic step to aid in SDS removal, minimizing residual irritative effects. In contrast to previous protocols that applied all reagents at once but achieved only partial decellularization and significant GAG loss(26), our controlled, sequential strategy ensures efficient cell removal while better preserving essential matrix components. This resulted in significant removal of cell-associated materials, with residual DNA levels below the recommended threshold (<50 ng/mg dry weight)(13). Similar to other methods, we report an impact on the deposited ECM principally in the form of a significant GAGs content reduction(22, 27). Nonetheless, changes did not impair the grafts bone-forming capacity in both stringent ectopic and orthotopic models. These findings highlight the ability of the sole extracellular matrix, independent of living cells, to drive endochondral ossification and support robust bone regeneration.

Despite the proven decellularization, robust safety assessment is essential before considering clinical translation. This is particularly important also in regard to the exploitation of a human cell line for the graft engineering process. Ectopic implantation in ID and IC animals are considered standard assays for teratoma and immunogenicity assessment. The ectopic implantation also presents the advantage of assessing active cell recruitment, as compared to extensive bleeding or inflammation typically resulting from orthotopic surgery. Our method allowed compiling a comprehensive temporal and quantitative analysis of immune recruitment, summarized in the form of immune prints. We evidenced the importance of the early innate invasion, dominated by macrophages recruitment similarly to early phases of bone repair(28). Comparing collected profiles in ID and IC, we could correlate the transition of macrophages from an M0 to M2 phenotype during the early inflammatory phase with subsequent success of ectopic ossification. This shift is consistent with the well-established sequence of immune events, where early proinflammatory responses are gradually replaced by a reparative phase characterized by the resolution of inflammation and tissue remodelling(1, 29, 30). M2 macrophages have been largely described as active promoter of angiogenesis, tissue repair, and associated with pro-regenerative processes in various tissue contexts including bone formation(1, 31–33). While informative, one limitation of our methodology relates to the simplified binary M1/M2 phenotypic definition, typically challenged by the tissue location and inflammation contexts(34, 35).

The ectopic model further allows comparing the in vivo immunogenicity of HyC pre- and post-decellularization. Surprisingly, the decellularization was shown to mostly affect the extent of recruited immune cells, but not their inflammatory profiles. As such, D-Hyc exhibited a reduced blood infiltration as compared to L-HyC, consistent across timepoints and animal models. Beyond immune recruitment, L-HyC and D-Hyc displayed very similar immune prints, as well as identical pro-inflammatory M1 versus pro-regenerative M2 ratios. Corroborated by the absence of an early M2 polarization, none of the grafts initiated an effective remodelling into bone upon implantation in IC mice. Of note, the apparent immune rejection was not associated with T cell expansion, whose frequency remained stable over time for both L-HyC and D-HyC. Existing studies have assessed the performance of living human cartilage in IC models(16, 36), relying on the immunosuppressive properties of differentiated BM-MSCs to circumvent immune rejection. However, effective bone formation could not be achieved in this xenogeneic setting. Since our D-HyC grafts are devoid of human cells, our study identifies interspecies variations in ECM proteins as the primary cause of human ectopic HyC rejection. This finding highlights the challenge of using animal models to co-jointly evaluate the immunogenicity and osteoinductivity of decellularized human grafts.

To complement ectopic findings, we thus established in vitro assays as simplified but fully human platforms mimicking an allogeneic response. While several studies evaluated the immunogenicity of living chondrogenic tissue(37),(38, 39) and BM-MSCs *in vitro*(40, 41), the inflammatory response induced by human decellularized grafts has not been reported. By exposing HyC to the full spectrum of antigen presenting cells, we could demonstrate that engineered extracellular matrix do not elicit a strong inflammation. Using multiple healthy donors, PBMCs and macrophages resulted in minimum T-cell activation for both CD4 and CD8 subsets, in line with low detection of pro-inflammatory cytokines. In addition to immuno-evasion, our engineered ECM grafts were also shown to impact the in vitro maturation/function of human immune cells. In particular, D-HyC significantly reduced both the M1 (pro-inflammatory) and M2 (pro-regenerative) macrophages polarization. Grafts were further demonstrated to affect DCs maturation, as evidenced by low levels of standard activation markers (CD80 and HLA-DR). These findings echo the previous results on chondrogenically differentiated hMSCs failing at inducing T-cell-activation and DC maturation(38, 39, 42, 43). However, our study extends previous findings by demonstrating that decellularized constructs, devoid of living cells, similarly exhibit comparable immunomodulatory properties. This is unexpected, as implying that our engineered ECM and embedded factors are sufficient to confer and retain potent immuno-privileged features, despite the lyophilization and decellularization processes. Taken together, findings from human in vitro assays strongly support the safety and allogeneic exploitation of D-HyC as a bone graft substitute with tolerogenic properties.

In a final step, D-HyC grafts were evaluated in a rat orthotopic model featuring a critical-sized femoral defect. Compared to pre-existing approaches, the graft demonstrated remarkable performance in terms of both reproducibility and bone repair efficacy. Several studies have explored the endochondral ossification pathway and tested HyC potential in orthotopic settings. To the best of our knowledge, our graft outperforms previous strategies by achieving faster and superior bone formation(6, 16, 18, 44–46). However, implantation of a living rat cartilage graft and high doses of BMP-2 also resulted in comparable healing at 12-weeks. Those strategies actually mimic current clinical gold standards, consisting in autologous grafting and BMP-2 delivery. Their limitations are well-known, reinforcing the relevance of our D-HyC, approach superior in safety and performance. Most importantly, unlike prior studies relying on living implants, our D-HyC is strictly off-the-shelf, and its regenerative performance was assessed without any additional materials. Beyond femoral bridging, mechanical restoration was also observed, although it did not reach the strength of native femurs. We hypothesize that the stabilizing plate may have reduced load-bearing and mechano-stimulation in the defect leg, thereby preventing full restoration of native mechanical properties.

Importantly, in sharp contrast to ectopic observations, the rat IC orthotopic environment did not lead to graft rejection. This highlights that beyond graft composition, the local environment plays a pivotal role in regenerative decision-making(47–49). More generally, the discrepancy across animal models and site of implantations challenge the establishment of standard criteria to evaluate safety and efficacy of human decellularized biomaterials. Their translation into clinical practice is further hindered by the lack of regulatory uniformity, as their classification straddles the line between cell-based and acellular products. This is particularly the case for products that do not involve a recellularization step with patient cells before implantation, such as our D-HyC. Recent regulatory advancements include the FDA approval of an engineered acellular vessel(50), resulting from primary human endothelial cells 3D differentiation and subsequently decellularized(51). Our D-HyC follows a similar approach, albeit exploiting a human immortalized line. This evidence a regulatory path for cell-based but cell-free products, here falling in the Regenerative Medicine Advanced Therapy Designation. A clear equivalent remains ambiguous under the European Medical Agency (EMA) classification, whereby the Tissue Engineered Product category appears the most relevant but has so far only been applied to living substitutes. A clear and uniform classification can only facilitate the journey to clinical translation, together with defining preclinical models and in vitro systems modeling the complexity of the human immune system and physiological responses(12, 52). This remains an open challenge, with performance in humans not being predictable particularly regarding their safety, integration, and longevity within the body.

## Conclusion

The development of decellularized tissue engineered grafts represents a promising approach for tissue regeneration, offering a scalable, off-the-shelf solution that bypasses the limitations of autologous and living grafts(19, 53). Using a dedicated human cell line, our work demonstrates the possibility to engineer D-HyC as bone graft substitute exhibiting unprecedent bone formation capacity together with intrinsic immunoregulatory functions. Additionally, it underscores the limitations of current in vitro and in vivo models in accurately assessing the immunogenicity and performance of decellularized human grafts.

## Materials and Methods

### MSOD-B culture

MSOD-B cells (P21) are expanded in T175 flasks in complete medium consisting of alpha Minimum Essential Medium supplemented with 10% fetal bovine serum, 1% HEPES (1M), 1% Sodium pyruvate (100mM), 1% of Penicillin/Streptomycin/Glutamine solution and 5ng/ml of FGFb (all from Gibco Invitrogen, USA) under standard culture conditions until confluence (90%). Medium is replaced twice a week. Cells are seeded at a standard density of 3.400 cells/cm2.

### MSOD-B seeding

MSOD-B cells (P23) are harvested and seeded on cylindrical type I collagen sponges (AviteneTM UltrafoamTM Collagen Sponge, Davol) of 6mm in diameter and 3mm in thick. Briefly, scaffolds are shaped with a 6mm biopsy punch (Kai Biomedical) and placed in a single well of a 12-well plate coated with 1% agarose to prevent cell adhesion. Per scaffold, 35µL of cell suspension containing 2M of cells are seeded at the surface and incubated 1h under standard culture conditions (cell culture media as seen before without FGFb). Then 2mL of chondrogenic medium is added to the well. This medium consisted of DMEM supplemented with 1% Insulin/Transferrin/Selenium, 1% Sodium pyruvate (100mM), 1% of Penicillin/Streptomycin/Glutamine (100X) solution (all from Gibco Invitrogen, USA), 0.47mg/mL linoleic acid (Sigma Aldrich, USA), 25mg/mL bovine serum albumin (Sigma Aldrich A2153-50G), 0,1mM ascorbic acid-2-phosphate, 10ng /ml TGFb3 and 10-7M dexamethasone. Chondrogenic medium is changed twice a week for a period of three weeks.

### Lyophilization

After three weeks of hypertrophic differentiation, MSOD-B constructs are rinsed twice with PBS 7,2 (Gibco Invitrogen, USA). For lyophilization, cell constructs are then snap froze inside 15-ml tubes in liquid nitrogen for 5 minutes and then lyophilized at -80°C and 0.05mbar over night. Once freeze dried, the lyophilized cell constructs (L-HyC) are stored at 4°C for long term storage.

### Decellularization

After lyophilization samples were immersed in a hypertonic solution (Tris-HCl 50mM, NaCl 1.5M, pH 7.6) for 10H at RT over continuous stirring (100rpm). After rinsing 3X with PBS at RT (30min each wash) each scaffold was immersed in a hypotonic solution (Tris-HCl 10mM, pH 8) ON at RT and over continuous stirring at 100rpm. Then samples were treated with SDS 0.5% in the hypotonic solution for 12H (RT, 100rpm). Once treated with the detergent, samples were washed ON with the hypertonic solution (RT, 100rpm), subsequently washed three times with PBS as before for a total duration of 6H and treated with DNase I 50u/mL (Tris-HCl 10mM, pH. 7,6) and Aprotinin 10KIU/mL for 3H at 37°C under continuous stirring (100rpm) then washed 3X with PBS at RT (30min each wash). Once decellularized samples were lyophilized prior to implantation and quantitative analysis.

### Powder formation

Only lyophilized (L-HyC) or decellularized and lyophilized (D-HyC) cartilages are then grinded with the CryoMill (RETSCH) in a 10 ml grinding jar of stainless steel with two 10 mm grinding balls. In order to ensure that the sample is pre-embrittled before grinding a pre-cooling time of 12 minutes is performed. Grinding is then performed for 4x 2 minutes at 25 hz and an intermediate cooling time of 30 seconds at 5Hz. Both powders are stored at 4°C for long term storage.

### Biochemical analysis

After lyophilization, engineered human cartilage grafts are digested overnight at 56°C in 0.5 mL of a Proteinase K solution (Proteinase K, 1 mg/mL; pepstatin A, 10 μg/mL; 1 mM EDTA; 100 mM iodoacetamide; 50 mM Tris, all from Sigma) at pH 7.6. DNA content was evaluated fluorometrically using the CyQUANT NF Cell Proliferation Assay Kit (Thermo Fisher, USA), following the manufacturer’s protocol (excitation at 485 nm, emission at 535 nm). Glycosaminoglycans (GAGs) content was analyzed by Glycosaminoglycan Assay Blyscan kit (Biocolor) following the manufacturer instruction. Total collagen was assessed by determining hydroxyproline content after acid hydrolysis (HCL 6N, 12H, 120°C), followed by a reaction with p-dimethylaminobenzaldehyde and chloramine T (Sigma-Aldrich, USA), applying a hydroxyproline-to-collagen ratio of 0.134. Both GAG and collagen contents are normalized to the dry weight of each sample. For BMP-2 protein content, human cartilage grafts were immersed in a RIPA buffer supplemented with protease inhibitor. Tissues are then lysed with a tissue homogenizer (TissueRuptor II, Qiagen) and total BMP-2 content within tissues was assessed using the human BMP-2 DuoSet ELISA (R&D Systems) according to the manufacturer’s instructions.

### Immunofluorescence staining

Samples were fixed in 4% paraformaldehyde (Thermo Scientific) at 4°C, for 24 h and decalcified with 10% EDTA solution (Sigma), pH 8 up to 2 weeks with gentle shaking, at 4°C. Embedding was performed using 4% low-melting agarose (Sigma) and 100 µm thick sections were cut using a 7000smz Vibratome (Campden) with stainless steel or ceramic blades (Campden). All staining and washing steps were performed under gentle shaking. Sections were blocked and permeabilized with 0.5% Triton X-100 (Sigma) in PBS and 20% donkey serum (Jackson ImmunoResearch) for 1 h, at room temperature (RT). After blocking/permeabilization, sections were stained with primary antibodies at 4°C, overnight, and highly cross-absorbed secondary antibodies for 2-3 h, at RT with 3 times 20 min washes in between using 0.1% Triton X-100 in PBS and 2% donkey serum. Tissue sections were treated with Vector TrueView Autofluorescence Quenching Kit (Vector Laboratories) to remove unwanted fluorescence. Tissue sections were mounted with a Vectashield antifade mounting medium containing DAPI (Vector Laboratories, H1200). Samples were imaged with an LSM780 confocal microscope (Zeiss). We used the following primary antibodies: mouse anti-Collagen II (Invitrogen, MA137493) and rabbit anti-Collagen Type X (COL10) (abbexa, abx101469. The following secondary antibodies were employed: CF633 donkey anti-mouse (Sigma-Aldrich, SAB4600131) and CF568 donkey anti-rabbit (Biotium, 20098).

### PBMCs preparation

Peripheral blood mononuclear cells (PBMCs) were isolated from heparinised blood using density gradient centrifugation (Lymphoprep, Axis-Shield) at 640g for 20 minutes with low break. The PBMCs were washed twice with PBS, counted, and prepared for subsequent assays. When required, monocytes were further isolated from PBMCs using CD14+ magnetic bead separation (Miltenyi) following the manufacturer’s instructions, counted, and used in downstream assays. CD4^+^ T-cells were isolated from the PBMC fraction using the EasySep™ CD4^+^ T-cell isolation kit (Stemcell Technologies) as per the manufacturer’s instructions.

### Monocyte isolation and polarization in vitro

Monocytes were isolated from freshly isolated PBMCs, as described above, from healthy controls upon informed consent (n=6, median age 43, 50% female). Monocytes were next cultured for 5-days in 24-well plates (Falcon) at 0.5x10^6^cells/ml in complete alpha Minimum Essential Medium supplemented with 10% heat-inactivated fetal bovine serum, 1% HEPES (1M), 1% Sodium pyruvate (100mM), 1% of Penicillin/Streptomycin/Glutamine solution and 40ng/ml M-CSF. After 5 days, media was replaced and supplemented with 200μg/ml powdered engineered human cartilage grafts (L-HyC or D-HyC) without M-CSF and cultured for an extra two days. As a control, both M1 and M2 macrophages were induced by stimulating monocytes with LPS (10 ng/mL) and IFN-γ (10 ng/mL) for M1 and IL-10 (25 ng/mL) + dexamethasone (10 nM) for M2.

### Macrophage polarization analysis and preparation for T-cell proliferation

Forty-eight hour in vitro polarized monocytes were detached using ice-cold PBS/1mM EDTA and gentle pipetting. Next, they were washed with PBS, counted and resuspended in RPMI-1640 supplemented with 2mM L-glutamine, 10% heat-inactivated fetal bovine serum, 1% of Penicillin/Streptomycin solution. 2.5x10^3^ cells were removed and used for the T-cell proliferation assay (see below). The remaining cells are washed with PBS and stained with anti-CD80, CD86 (clone FUN-1, BV650, BD), CD163 (clone: GHI/61, PE, BD) and CD206 (clone:19.2, APC/Fire750, BD), all diluted 1:100, for 20min, RT. Finally, they were washed once more with PBS and analysed by flow cytometry (CytoFLEX).

### T-cell proliferation assay

T-cells isolated as described above were stained with 2µM CellTrace Violet (Invitrogen) for 20min at 37°C. Remaining dye was quenched by the addition of 6X the volume of complete medium. The cells were centrifuged, resuspended in complete medium and counted. A 96-well plate (Eppendorf) was coated with anti-CD3 (1:1000, Clone OKT3, Invitrogen) for 90min. The coating solution was removed prior to use. Wells without coating served as negative controls. In addition, anti-CD28 (clone CD28.2, Invitrogen, Thermo Fisher Scientific, 1:1000) coating was added in some wells as a positive control. Macrophages from the in *vitro* polarization assay were counted (XN-350, Sysmex) and resuspended in RPMI-1640 supplemented with 10% foetal calf serum, 2mM L-glutamine and PenStrep. Next, macrophages and T-cells at a 1:20 ratio (monocytes/macrophages: T-cells) were added to the coated plate in a total volume of 200µl. The cells were incubated for 72h, 37°C, at 5% CO_2_. Subsequently, the cells were detached through gentle pipetting, centrifuged, and stained with anti-CD3 (clone UCHT1, alexa fluor 700, 1:200), anti-CD25 (clone M-A251, PerCP-Cy5.5, 1:200), anti-HLA-DR (clone G46-6, APC-H71:200) and anti-CTLA-4 (clone BNI3, PE, 1:50), all from BD, for 20min, RT. Finally, the cells were washed once with PBS and analysed using flow cytometry (CytoFLEX).

### PBMCs analysis following powder activation

Previously lymphoprep isolated PBMCs were stained with 2µM CellTrace Violet (Invitrogen) as above. PBMCs were then resuspended in RPMI-1640 supplemented with 10% foetal calf serum, 2mM L-glutamine and PenStrep and culture at 0.5x10^6^ in 48-well plates (Falcon). Phytohemagglutinin-M (PHA-M, 10µg/ml, Roche) was added in some wells as a positive control. PBMCs were then supplemented with 200μg/ml powdered engineered human cartilage grafts (L-HyC or D-HyC) and cultured for five days. At the end of the culture, media was collected for supplementary cytokines analysis and the cells were detached through gentle pipetting, centrifuged, and stained with anti-CD3 (clone UCHT1, alexa fluor 700, 1:200), anti-CD4 (clone RPA-T4, alexa fluor 488, 1.5:100), anti-CD8 (clone RPA-T8, PeCy7, 1.5:100), anti-CD19 (clone SJ25-C1, BV786, 1:200), anti-CD25 (clone M-A251, PerCP Cy5.5, 1:200), anti-HLA-DR (clone G46-6, APC-Cy7,1:200), all from BD, for 20min, RT. Finally, the cells were washed once with PBS and analysed using flow cytometry (CytoFLEX). IL-6, IL-10 and IFNg content within the supernatant was assessed using DuoSet ELISA (R&D systems) according to the manufactureŕs instructions.

### Monocyte-derived dendritic cells (Mo-DCs) preparation and culture

Monocytes were isolated from freshly isolated PBMCs, as described above. Freshly isolated monocytes were seeded at 1x10^6^cells/ml in 24-well plate and cultured for five days following the ImmunoCult^TM^ Dendritic Cell Culture Kit (STEMCELL) instructions. After five days, unmatured Mo-DCs were supplemented with 200μg/ml powdered engineered human cartilage grafts (L-HyC or D-HyC), ImmunoCult^TM^ Dendritic Cell Maturation Supplement (STEMCELL) or left untreated for 48h, 37°C, at 5% CO_2_. At the end of the culture, cells were detached using ice-cold PBS/1mM EDTA and gentle pipetting. Next, they were washed with PBS, counted and stained with anti-CD14 (clone 63D3, FITC, BioLegend, 1:100), anti-CD80 (clone 2D10, BV650, 1:100, BioLegend), anti-HLA-DR (Clone G46-6, PE-Cy7, BD, 1:100).

### Ectopic implantation

*FoxN1* KO BALB/C (Nude mice) & C57BL/6J (wild type) mice of 6–8-week-old were obtained from Charles River Laboratories. All mouse experiments and animal care were performed in accordance with the Lund University Animal Ethical Committee (M15485–18) under the regulation of the Swedish board of agriculture following the 3R’s principles. Mice were housed at a 12-hour light cycle in individually ventilated cages at a positive air pressure and constant temperature. Mice were fed with rodent chow and sterile water. Immune recruitment, tissue remodeling and bone formation efficiency of engineered human cartilage grafts is assessed by ectopic subcutaneous pouches in both ATHYM-Foxn1nu/nu mice and C57BL/6J mice, with a maximum of 6 implants per animal. For surgical procedures, animals are anesthetized by inhalation using a mixture of oxygen (0,6mL min-1) and isoflurane (1.5-3vol%). After -3, -5 and-7-days or 6- and 12-weeks post implantation, samples were explanted and fixed in 4% formaldehyde solution (Solveco AB, Sweden) ON at 4°C prior to microtomography and histological analysis (Safranin-O staining) or immediately digested for FACS analysis.

### Critical-sized femoral defect implantation

Ten- to 12-week-old male Sprague Dawley rats (n=15) were purchased from JANVIER LABS (France). After 7 days of acclimation the rats were anesthetized using 3% isoflurane. Then, animals were placed on a 37°C warm heating pad in prone position. Once anesthetized, isoflurane was lowered to 2 to 2.5%, and buprenorphine (Temgesic, 30 μg/kg; Indivior Europe Ltd., Dublin, Ireland) was injected subcutaneously for analgesia. The right hind limb of the animal was shaved and carefully disinfected, and an incision was made along the skin and soft tissue to expose the right femur. After cleaning the femur laterally from soft tissue, a customized in house-developed four-hole internal fixation plate (Ø of 1.5-mm straight locking plate, PEEK) was held to the lateral aspect of the femur using forceps and a clamp. The plate was then screwed to the bone using two proximal and two distal screws (Ø of 1.5-mm locking screws, stainless steel; outer screws, 7 mm in length; inner screws, 6 mm in length; DePuy Synthes). After fixation, a 5-mm osteotomy was performed using two Gigli wires (0.44 mm; RISystem AG, Landquart, Switzerland) with the help of a custom-made, three-dimensional (3D) printed saw guide. The defect was then filled with three decellularized grafts. The grafts were push-fitted and not secured with any sutures. The wound was closed in a layered fashion using resorbable sutures (Vicryl 4-0, Ethicon, Somerville, USA) by closing the muscle first (continuous interlaced suture), followed by closing the skin (Donati suture). Animals started load bearing immediately after surgery. After 6 (n=2) or 12 weeks (n=15) animals were anesthetized by isoflurane inhalation (3%), followed by CO2 asphyxiation, and right-femur were harvested prior to subsequent micro-CT, mechanical (n=7) and histological (n=3) characterization. The control empty group used in this study was part of a parallel experiment and has been previously published(17). The animals, surgical procedures, and post-operative care for this group were performed under identical conditions and protocols as the current study to ensure comparability.

All rat experiments and animal care were performed in accordance with the Swedish Board of Agriculture approval (permit number: 18-08106/2018) following the 3R’s principles.

### In vivo Micro-CT scanning

All rats were subjected to *in vivo* x-ray analysis at 4-6 weeks after surgery. Briefly, animals were anesthetized using isoflurane (3%) and placed in a right lateral decubitus position. Lower body was then scanned with a U-CT system (MILABS, Netherland) using a tungsten x-ray source at 50 kV and 0.21mA. A circular scan (360°) was recorded with an incremental step size of 0.250°. Volumes were reconstituted at 30µm isotropic voxel size using MILABS software analysis. For bone volume analysis, the highly mineralized tissue volume was quantified using Seg3D (v2.2.1, NIH, NCRR, Science Computing and Imaging Institute (SCI), University of Utah, USA). For total volume analysis, each sample was meshed with Blender (v2.82a, Netherland) and analyzed with an in-house developed script.

### Ex vivo Micro-CT scanning

Subcutaneous (mice) and orthotopic (rat) implanted samples were fixed overnight with 10% formaldehyde before being subjected to ex-vivo micro-CT with a U-CT system (MILABS, Netherland) using a tungsten x-ray source at 50 kV and 0.21mA for subcutaneous retrieved samples and 65KV and 0.13mA for rat-femurs. Volumes were reconstituted at 10µm isotropic voxel size and analyzed for bone volume and total volume as described in the previous section.

### Mechanical testing

Femurs bearing defects and contralateral femurs were subjected to 3-point bending following previously established protocol(17). Briefly, specimens were positioned on a 3-point bending jig with 16 mm support spacing, loaded in the antero-posterior position, ensuring the defect was centred. Testing was performed using an Instron 8511.20 load frame, applying a 20 N pre-load for 10 s, followed by axial compression at 0.25 mm/s until fracture. Peak force and stiffness (linear slope) were extracted from the force-displacement curve.

### Histological staining

L-HyC, D-HyC both in vitro or in vivo were washed in 1x PBS after formaldehyde fixation. Explanted in vivo tissues were decalcified with 10% EDTA (Sigma Aldrich, USA), pH 8.0, at 4°C for 2 weeks before paraffin-embedding. Tissues were progressively dehydrated using graded ethanol solutions (35%, 70%, 95%, and 99.5%; Solveco), with two 20-minute immersions per concentration. Following dehydration, the tissues were washed in a 1:1 mixture of 99.5% ethanol and xylene (Fisher Scientific) for 10 minutes, then treated with xylene alone for 20 minutes, twice. Subsequently, the tissues were embedded in paraffin at 56°C overnight and sectioned into 7–10 μm slices using a microtome. These sections were dried at 37°C overnight. To deparaffinize, the sections were rinsed twice in xylene for 7 minutes each, followed by a single 3-minute wash in 1:1 ethanol/xylene (99.5%). Rehydration was then performed using a graded ethanol series (99.5%, 95%, 70%, and 35%), with each step lasting 7 minutes and repeated twice.

### Safranin-O staining

Sections Sections were stained with Mayer’s hematoxylin solution (Sigma-Aldrich) for 10 minutes, followed by a rinse in distilled water to eliminate excess stain. Next, they were treated with 0.01% fast green solution (Fisher Scientific, USA) for 8 minutes, and any surplus dye was quickly removed by rinsing the sections in 1% acetic acid solution (Sigma Aldrich, USA) for 15 seconds. The slides were then stained with 0.1% safranin O solution (Fisher Scientific, USA) for 8 minutes. Dehydration and clearing were carried out by immersing the slides sequentially in 95% and 99.5% ethanol, followed by a 1:1 mixture of 99.5% ethanol and xylene. Finally, the sections were washed twice in xylene for 2 minutes to remove any residual ethanol and mounted using PERTEX mounting medium (PERTEX, HistoLab).

### Massońs trichrome staining

Trichrome staining was conducted using the Trichrome Staining Kit (Sigma-Aldrich) in accordance with the manufacturer’s protocol. Tissue sections were deparaffinized and rinsed as previously described. The sections were then placed in Bouin’s solution (Sigma-Aldrich) either overnight at room temperature or for 15 minutes at 56°C. Following this, the slides were washed under running tap water and stained with Weigert’s iron hematoxylin working solution (prepared by mixing equal volumes of solution A and B, EMD Millipore) for 5 minutes to visualize nuclei (stained black). After a wash step, the cytoplasm was stained red using Biebrich Scarlet-Acid Fuchsin for 5 minutes. The slides were then cleared by immersing them in a working phosphotungstic/phosphomolybdic acid solution (25 mL phosphotungstic acid, 25 mL phosphomolybdic acid, and 50 mL distilled water) for 5 minutes. Collagen fibers were stained blue by treating the slides with aniline blue solution for 5 minutes, followed by a clearing step in 1% acetic acid solution (prepared with glacial acetic acid, Fisher Scientific) for 2 minutes and a rinse under running deionized water. Finally, sections were dehydrated by sequential immersion in graded ethanol solutions (95% once, 100% twice) for 2 minutes each, followed by two 2-minute washes in xylene, and mounted with PERTEX mounting medium.

### Flow cytometry

The experiment on immune cell recruitment was carried out using flow cytometry (FACS). Under sterile conditions, harvested subcutaneously implanted samples were carefully cleaned of the connective tissue. Three samples were pooled together for a single FACS sample and digested in an enzyme cocktail consisting of 300 U/mg collagenase type II (Sigma-Aldrich, U.S.A), 2 U/mg collagenase P (Sigma-Aldrich, U.S.A) and 2 mM CaCl_2_ solution. The tissue digestion was carried out in a 12-well plate with 2 mL enzyme solution/well for 90 min in a humidified incubator at 37 °C. Any tissue remnants were mechanically homogenized using a 10 mL pipette by repeatedly passing the tissue through the pipette tip. The enzymatic reaction was stopped by mixing an equal volume of cell culture medium containing 10% (v/v) fetal calf serum to the digest. Then the solution was passed through a 70 µM tissue strainer and collected in FACS tubes (BD, U.S.A). The cells from the digested tissues were centrifuged at 1500 rpm for 5 min, the supernatant was discarded, and the cells were resuspended in 1 mL FACS buffer (PBS 2% v/v, 2 mM EDTA). 10 ul of sample was taken and mixed with equal volume of Tryphan blue and counted using Biorad TC20™ cell counting slides (Bio-Rad, Sweden). Cell suspension was re-spun to form a cell pellet, which was resuspended in 100 µL FACS buffer after which the respective primary antibodies for innate and adaptive immune system were added to the cell suspension and incubated at 4 °C for 1 h. Antibody dilutions were added to compensation beads for setting up fluorescence compensation (UltraComp eBeads™ compensation beads, Thermofisher Scientific). At the end of incubation, cells were centrifuged, and the supernatant was discarded. Cell pellets was washed with 1 mL FACS buffer, centrifuged, and re-suspended in 350 µL FACS buffer containing nuclear stain DAPI (500 ng/ul). The cells were then analyzed on a BD LSRFortessa™ cell analyzer (BD, U.S.A).

Antibodies list: APC Fire CD11b (Rat IgG2b, κ) (Biolegend 101235), PE F4/80 (Rat IgG2a, κ) (Biolegend 123110), PECy7 CD11c (Armenian Hamster IgG) (Biolegend 117318), AF647 CD206 (Rat IgG2a, κ) (Biolegend 141712), BV650 NK1.1 (Biolegend 108736), PerCPCy5.5 CD45 (Rat IgG2b, κ) (Biolegend 147706), CD16/32 (Rat IgG2a, λ) (Biolegend 101302), APC Fire 750 CD8 (Rat IgG2b, κ) (Biolegend 344746), PE CD19 (Rat IgG2a, κ) (Biolegend 152408), PECy7 CD3 (Armenian Hamster IgG) (Biolegend 100220), AF647 CD4 (Rat IgG2b, κ) (Biolegend 100424).

## Acknowledgments

Lund University Bioimaging Centre (LBIC), Lund University, is gratefully acknowledged for providing experimental resources. The work was supported by the Knut and Alice Wallenberg Foundation, the Medical Faculty at Lund University, and Region Skåne. This project has received funding from the European Research Council (ERC) (Starting grant #948588 to P.E.B.)., from the Swedish Research Council (2019-01864 to P.E.B) and Sten K Johnsons Stiftelse

---

**Supp. Figure 1:**
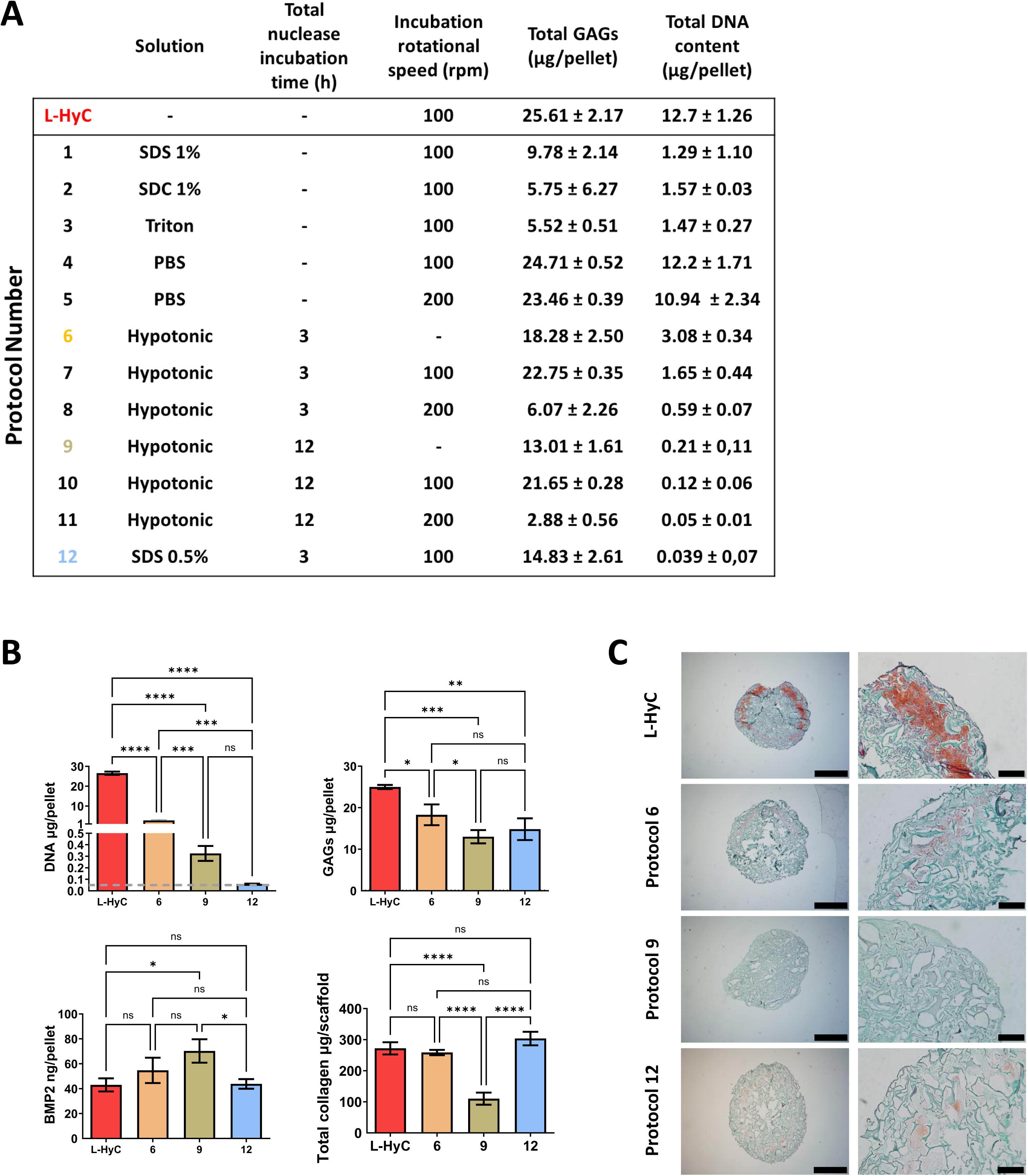
Development of a decellularization protocol. **A.** Protocol specification and resulting GAGs and DNA analysis performed on corresponding decellularized pellets (n≥3). **B.** Comparison of DNA, GAGs, Collagen and BMP-2 content on grafts after selected decellularization (n=6). Graphs represent mean ± standard deviation (SD), \**p* ≤ 0.05, \*\**p* ≤ 0.01, \*\*\**p* ≤ 0.001, \*\*\*\**p* ≤ 0.0001, determined by one-way ANOVA with Tukey’s multiple comparisons test. **C.** Representative Safranin-O staining images of pellets exposed to the indicated decellularization protocol (scale bar= 1mm and 200µm from left to right respectively).

**Supp. Figure 2:**
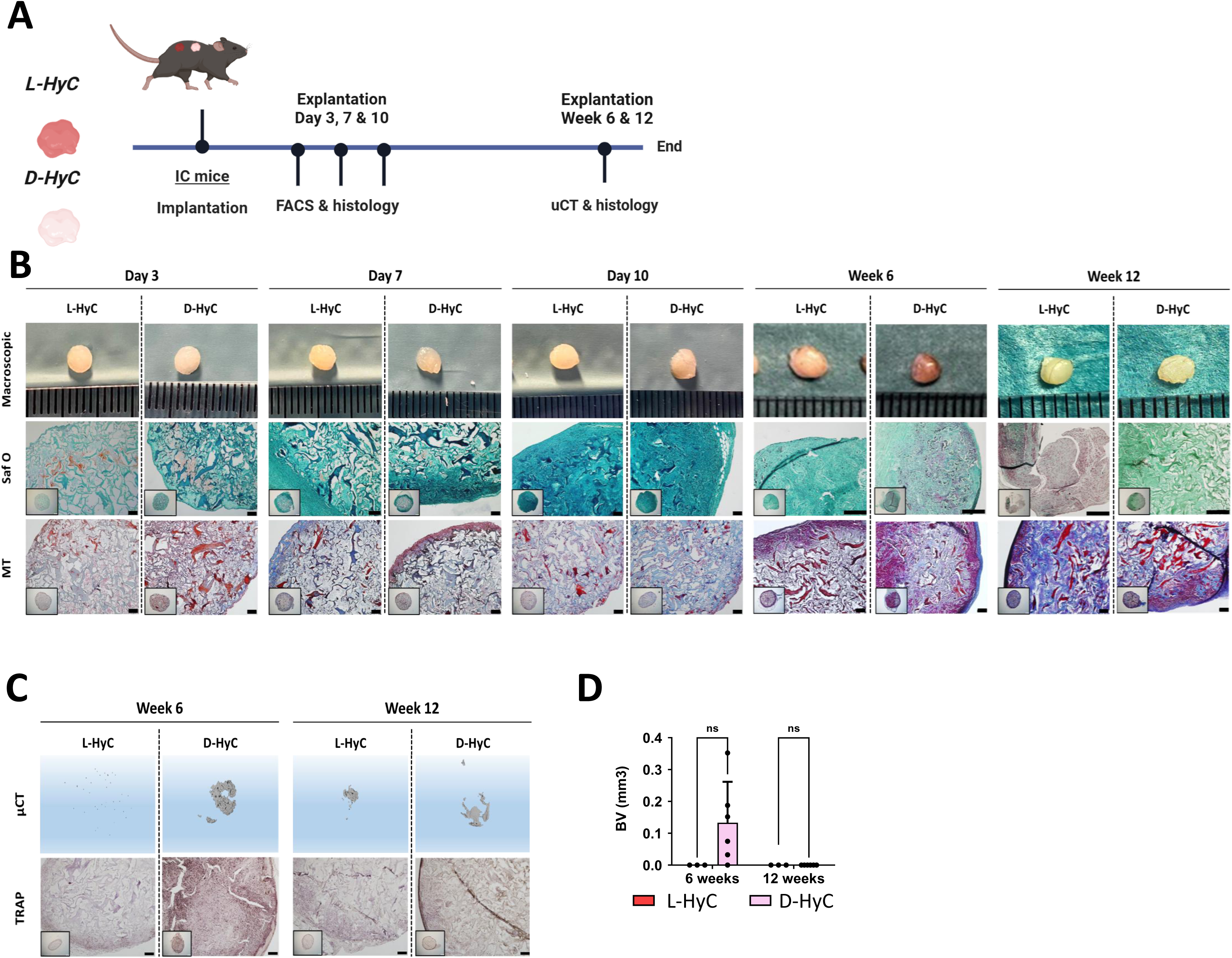
Decellularized human cartilage does not induce robust ectopic bone formation in immunocompetent mice. **A.** Experimental scheme for the *in vivo* ectopic osteogenic assessment of L-HyC and D-HyC grafts. **B.** From top to bottom, representative macroscopic images (scale bar=1mm), histological images of Safranin-O and Masson Trichrome staining of explanted tissues (Scale bar =100µm). **C.** From top to bottom, representative µCT 3D reconstruction and TRAP staining of explanted tissues at 6- and 12-weeks (Scale bar =400µm). **D.** BV/TV of µCT performed over explanted bones at 6- and 12-week post-implantation. The graphs represent mean ± standard deviation (SD), ns *p* > 0.05, determined by Two-way ANOVA.

**Supp. Figure 3:**
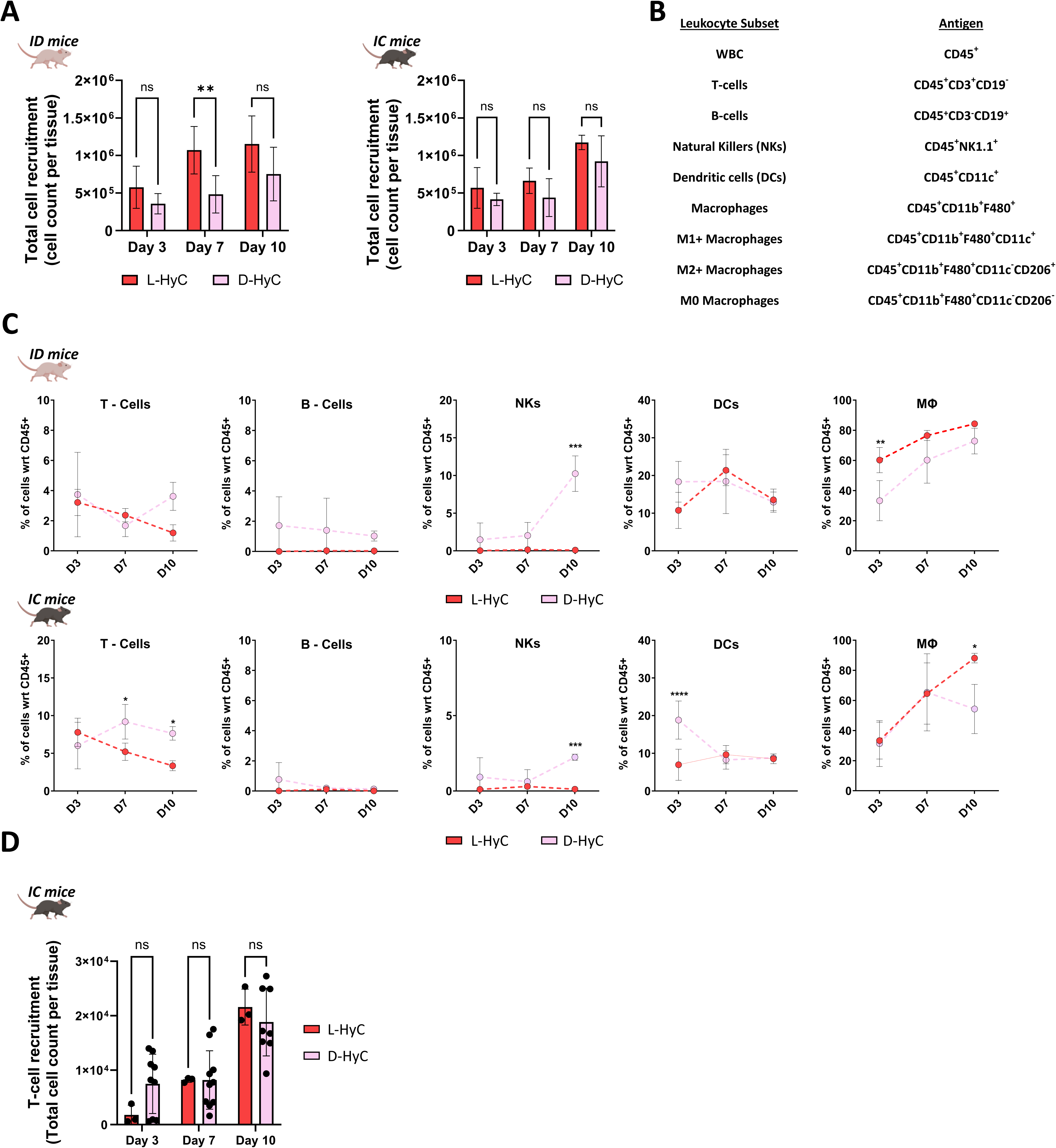
Immunogenicity assessment of engineered hypertrophic cartilage tissues in mice. **A.** Total cell number captured in explanted tissues after 3-, 7-, and 10-days post-implantation. The graphs represent mean ± standard deviation (SD), \**p* ≤ 0.05, \*\**p* ≤ 0.01, \*\*\**p* ≤ 0.001, \*\*\**\*p ≤ 0.0001*, determined by Two-way ANOVA. **B.** Flow cytometer gating strategy for T-cells, B-cells, NKs, DCs, and Macrophages. **C.** From right to left, percentages of early time recruitment of T-cells, B-cells, Natural Killers, Dendritic cells and Macrophages captured in explanted tissues after 3-, 7-, and 10-days respectively, in immunodeficient (Top) and immunocompetent (Bottom) animals. Error bars: The graphs represent mean ± standard deviation (SD), \**p* ≤ 0.05, \*\**p* ≤ 0.01, determined by Two-way ANOVA. N≥3 independent experiments, 3 pooled samples per animal (n ≥9). **D.** Total T-cell number captured in explanted tissues after 3-, 7-, and 10-days post-implantation in IC mice.

**Supp. Figure 4:**
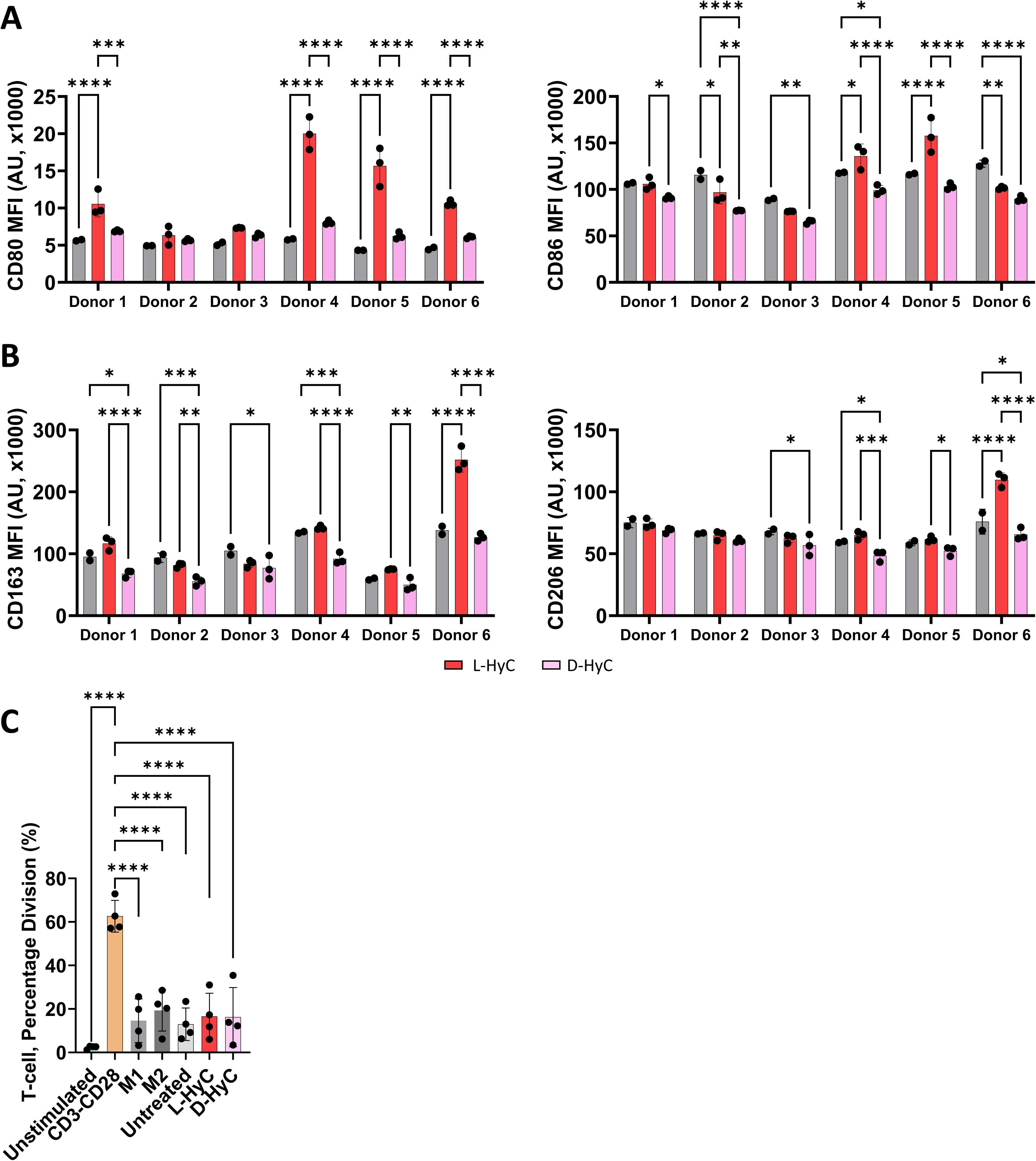
In vitro Immunogenic assessment of engineered hypertrophic cartilage tissues co-cultured with donor macrophages. **A.** Total CD80+ and CD86+ MFI indicating M1 polarization after 5-days co-culture with both L-HyC or D-HyC powdered cartilages (n=3 per donor). **B.** Total CD163+ and CD206+ MFI indicating M2 polarization after 5-days co-culture with either L-HyC or D-HyC powdered cartilages (n=3 per donor). The graphs represent mean ± standard deviation (SD), \**p* ≤ 0.1 \*\**p* ≤ 0.01, \*\*\**p* ≤ 0.001 \*\*\*\**p* ≤ 0.0001, determined by Two-way ANOVA. CD3+ percentage division assessed by FACS after co-culture of 5x10^4^ CD3+ cells with 2.5x10^3^ macrophages co-cultured with either L-HyC or D-HyC powdered cartilages. The graphs represent mean ± standard deviation (SD), *** *p* ≤ 0.001, \*\*\*\**p* ≤ 0.0001, determined by Ordinary one-way ANOVA.

**Supp. Figure 5:**
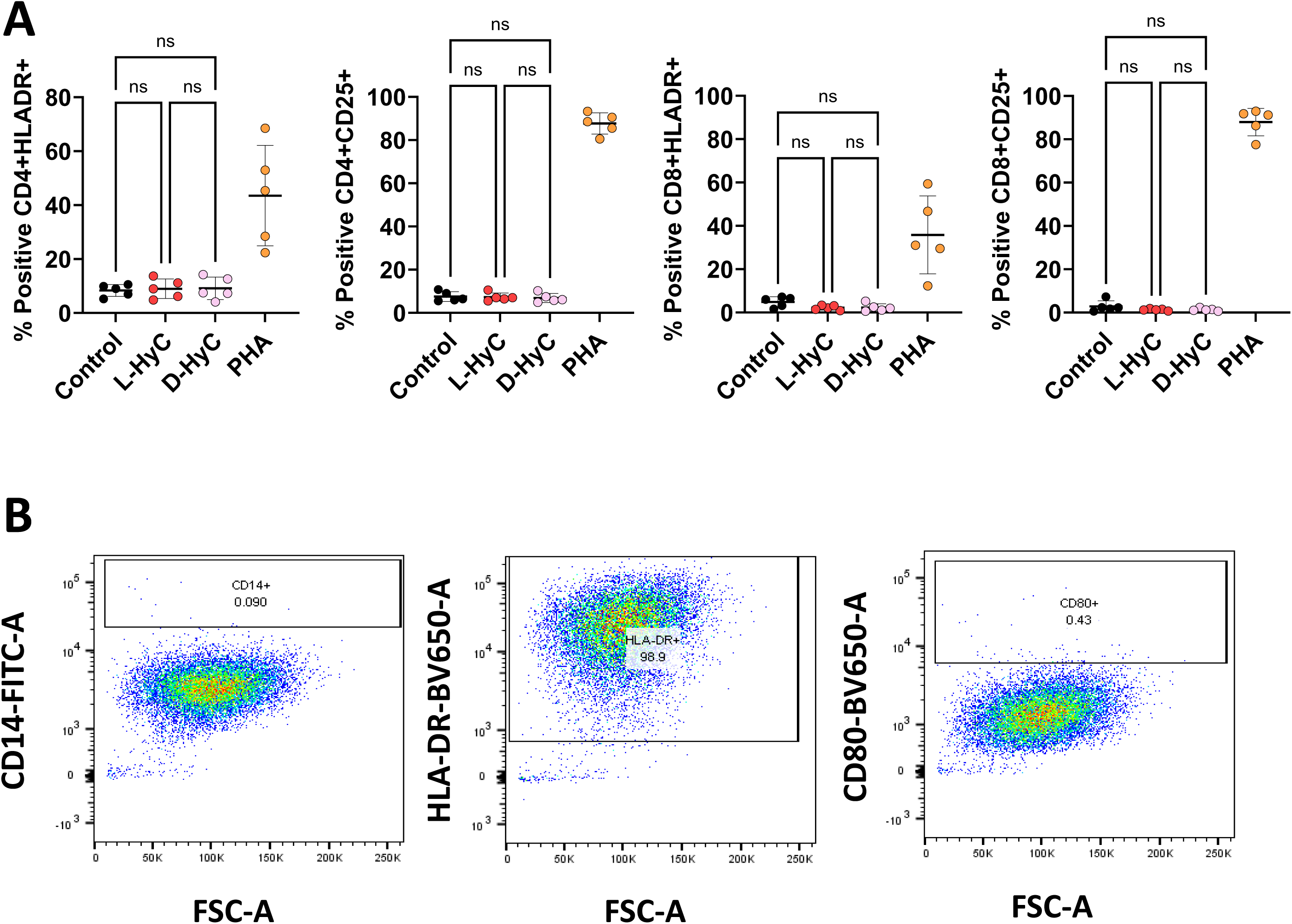
In vitro Immunogenic assessment of engineered hypertrophic cartilage tissues co-cultured with donor PBMCs or Mo-DCs. **A.** Percentage of CD3+/CD4+/HLA-DR+, CD3+/CD4+/CD25+, CD3+/CD8+/HLA-DR+ and CD3+/CD8+/CD25+ subtypes of T-cells derived from human donor PBMCs co-cultured with powdered L-HyC or D-HyC for 5-days analyzed by flow cytometry (5 donors, n=2 per donor). The graphs represent mean ± standard deviation (SD), determined by repeated measures one-way ANOVA, statistical significance set at *p < 0,05*. **B.** Representative FACS plots for monocytes derived dendritic cells (Mo-DCs) unstimulated with cartilaginous powder, indicating full dendritic differentiation (CD14-HLA-DR+) and lack of maturation (CD80+).

**Supp. Figure 6:**
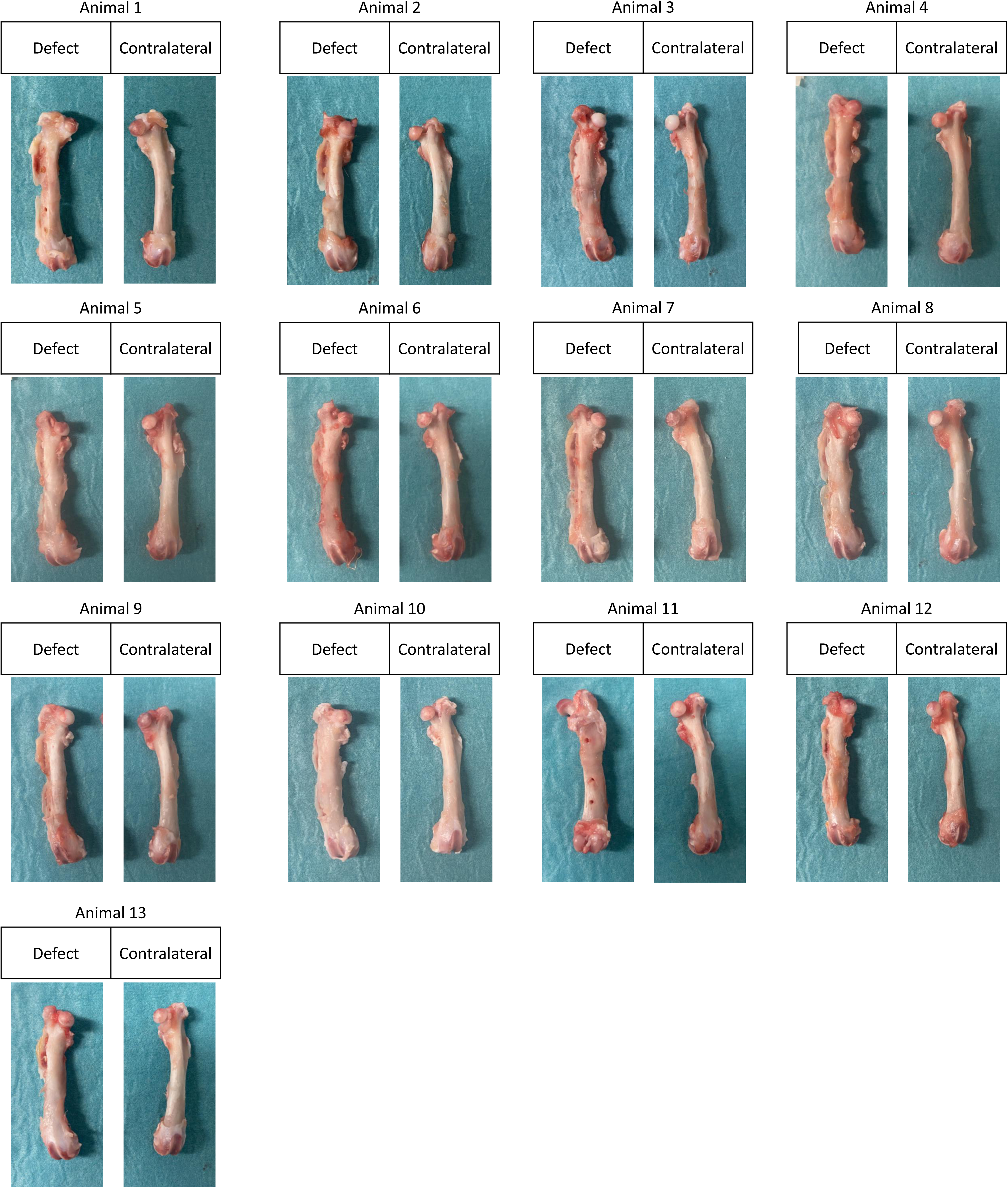
Macroscopic images of explanted femurs at 12 weeks post-implantation, showing the defect site (left) and contralateral control (right) for each animal.

**Supp. Figure 7:**
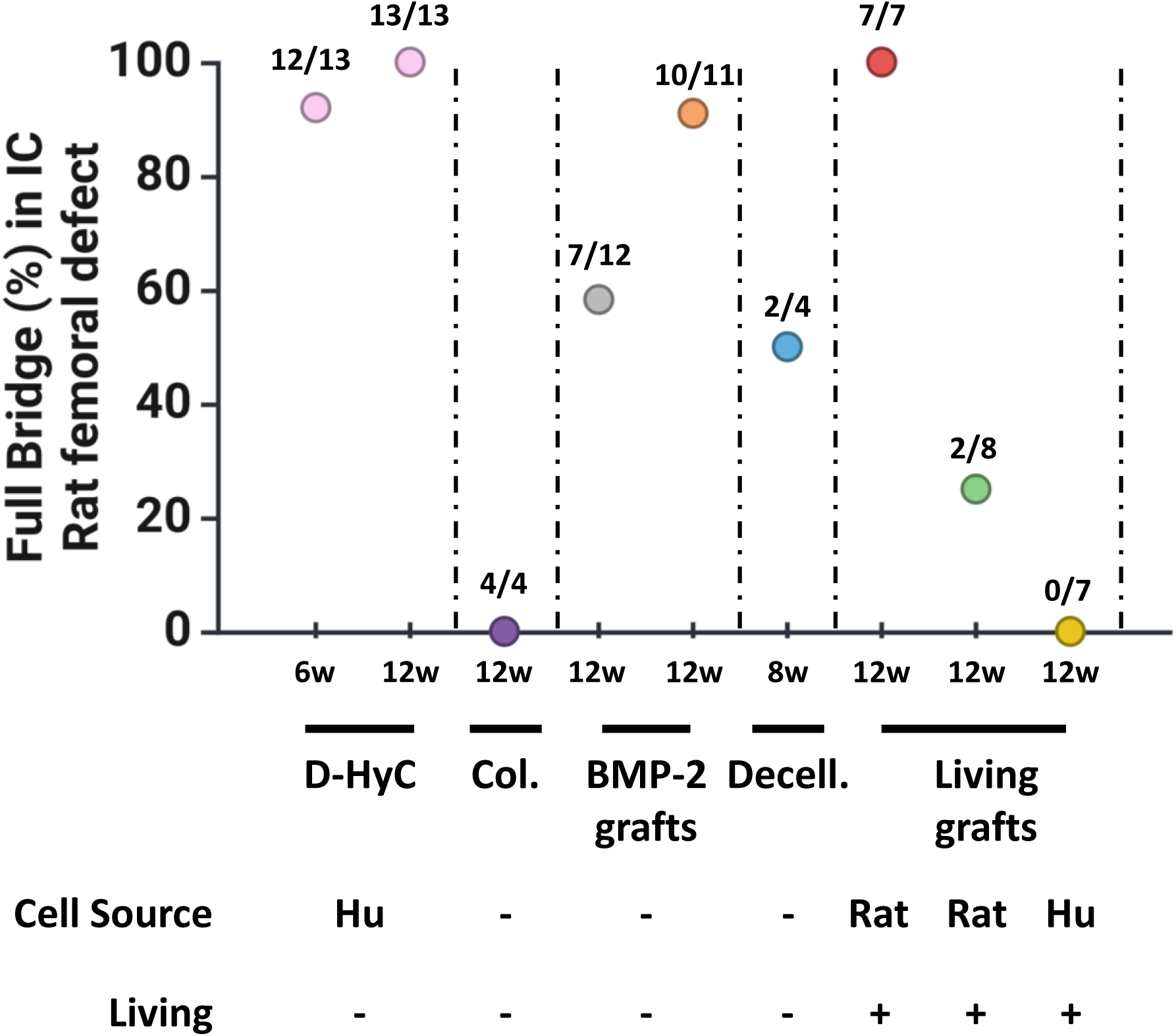
Performance comparison of different bone regeneration strategies in critical-sized femoral defects in immunocompetent rats. The graph shows the percentage of defects achieving full bone bridging at a specified time point for each graph (x-axis) with our **D-HyC** grafts (pink symbol). This comparison highlights the superior performance of **D-HyC** grafts, which achieved full bridging faster than all other strategies, including those requiring supraphysiological doses of BMP-2 (up to 10 µg per scaffold). • **Collagen scaffold (Col.)**: Collagen-only scaffold (purple symbol) used as a control group, data extracted from Longoni et al., 2020 (Ref. 16). • **BMP-2 grafts:** Collagen-based scaffolds loaded with either 0.5 µg (grey symbol, left) or 10 µg BMP-2 (orange symbol, right), from adapted from Liu et al., 2023 (Ref. 17). • **Decellularized scaffolds (Decell.):** Decellularized self-assembled human derived BM-MSC scaffolds (blue symbol) from Cunniffe et al., 2015 (Ref. 18). **-Living grafts:** Includes syngeneic (red symbol, left), allogeneic (green symbol, middle), and xenogeneic (yellow symbol, right) living tissue engineered cartilage grafts, adapted from Longoni et al., 2020 (Ref. 16). **-Cell Source** refers to the species origin of the graft material (Hu = Human, Rat = Rat), and **Living** indicates whether the graft contained viable cells at implantation.

